# Ultra-high-field fMRI reveals layer-specific responses in the human spinal cord

**DOI:** 10.1101/2025.07.17.665316

**Authors:** Ulrike Horn, S. Johanna Vannesjo, Nicolas Gross-Weege, Robert Trampel, Yulia Revina, Merve Kaptan, Alice Dabbagh, Laura Beghini, Virginie Callot, Andrew Todd, Kerrin J. Pine, Harald E. Möller, Jürgen Finsterbusch, Nikolaus Weiskopf, Falk Eippert

## Abstract

Developments in functional magnetic resonance imaging (fMRI) at ultra-high field (UHF) now allow for insights into human brain function at mesoscopic scale. However, similar progress has not been achieved for the human spinal cord, despite its importance for reciprocal brain-body communication. Here, we therefore integrate substantial methodological improvements that enable previously unattainable high-resolution UHF spinal fMRI. Using a heat-pain paradigm in healthy volunteers, we investigate fMRI responses in the dorsal horn: we characterize their spatial pattern, establish their robustness and demonstrate that they can be reliably observed even for individual participants. Most importantly, we uncover two physiologically relevant spinal response components – phasic and tonic heat responses – that map differentially onto superficial and deep layers of the dorsal horn. Taken together, our novel UHF fMRI approach allows for differentiating responses across fundamental spinal processing units and will enable insights into human spinal cord function at mesoscopic scale in health and disease.

## 1. Introduction

The spinal cord is a critical processing hub for bidirectional brain-body communication, relaying motor and autonomic output from, and somatosensory input to, the brain (Abraira & Ginty, 2013; Hochman, 2007; Lemon, 2008; Sengupta & Bagnall, 2023). At a macroscopic level, the spinal cord grey matter is organized into segments along the rostro-caudal axis and divided into dorsal and ventral horns segmentally; at the mesoscopic level, each horn contains fine-grained layers (Rexed, 1952, 1954; Silverdale, 2015; Tan et al., 2023). In animal models, a diverse array of sophisticated techniques is available to study spinal cord function at the macroscopic and mesoscopic level (Ahanonu et al., 2024; Claron et al., 2021; Shekhtmeyster, Carey, et al., 2023; Shekhtmeyster, Duarte, et al., 2023; Wu et al., 2019; Yang et al., 2015), allowing for insights at relevant spatial scales. Such insights also carry substantial translational importance, considering the spinal cord’s prominent role in sensorimotor disorders in humans (Ahuja et al., 2017; Ciccarelli et al., 2019; Kuner, 2015).

Human neuroscience however, lacks methods for investigating spinal cord function at the mesoscopic level. Although electrophysiological approaches (Spedden et al., 2024) allow for direct assessments of human spinal cord function with high temporal resolution, they fundamentally lack spatial precision. Functional magnetic resonance imaging (fMRI) does offer such spatially resolved recordings (Kinany et al., 2020; Nash et al., 2013; Oliva et al., 2022; Tinnermann et al., 2017; Vahdat et al., 2015, 2020), yet current spinal fMRI studies are mostly limited to magnetic field strengths of up to 3T. They are thus only able to deliver insights at the macroscopic spatial scale, leaving the fundamental functional units (layers) untouched and thereby preventing crucial neurobiological questions from being addressed.

This stands in sharp contrast to fMRI of the brain. There, high-resolution ultra-high-field (UHF) fMRI investigations at a magnetic field strength of 7T have for example delineated tonotopic maps in auditory cortex, orientation columns in primary visual cortex, and layer-specific input and feedback signals in somatosensory cortex (Formisano et al., 2003; Yacoub et al., 2008; Yu et al., 2019). Importantly, this provides access to fundamental computational units (for reviews, see Bandettini et al., 2021; De Martino et al., 2018; Uğurbil, 2021). Such access to relevant mesoscale elements is not currently possible in the spinal cord, neither at 3T nor with initial attempts at 7T (Barry et al., 2014; Conrad et al., 2018; Seifert et al., 2024). This leaves the need for high-resolution spinal data unmet, despite the human spinal cord being a prime target for UHF fMRI, considering its very small cross-section and intricate organization of grey matter into layers.

There are numerous spinal cord specific obstacles to overcome on the road to UHF fMRI (Barry et al., 2017), with no guarantee that techniques developed for 7T brain fMRI or 3T spinal fMRI are directly applicable. As an example, static B_0_ field inhomogeneities scale with field strength and are extensively present in the spinal cord at both small and large spatial scales (Cohen-Adad, 2017), leading to severe signal loss and geometric distortions with standard echo-planar-imaging (EPI) acquisitions. In this study, we addressed these and other longstanding challenges, with the goal of creating a spinal imaging approach with mesoscopic resolution. This would allow access to the fundamental computational units and thus offer a previously missing view on healthy and aberrant human spinal cord function.

To demonstrate the robustness of our strategy – which is based on numerous improvements in data acquisition and analysis – we acquired spinal cord UHF fMRI data at 7T from a large sample of healthy volunteers divided into screening, discovery, and preregistered validation datasets. Spinal cord blood-oxygen-level-dependent (BOLD) responses were evoked by individually calibrated painful heat stimuli, targeting the dorsal horn, i.e. the first synaptic relay in the central nervous system. We chose the domain of pain as a test-bed for our approach due to three reasons. First, to ensure neurobiological relevance, since the spinal cord is the initial bottom-up processing site for nociceptive stimuli in the central nervous system and also the target of powerful top-down control mechanisms (Fields, 2004; Heinricher et al., 2009). Second, to demonstrate translational applicability, as spinal mechanisms are extensively involved in pain chronification (Colloca et al., 2017; Kuner, 2015). Finally, to guarantee the generalizability of our approach beyond the domain of pain, because the dorsal horns (housing sensory functions) are much thinner and thus more difficult to investigate than the ventral horns (housing motor functions).

## 2. Results

### 2.1. Improving data acquisition and analysis methodology

Our approach for data acquisition involved several methodological developments that allowed for the acquisition of high-quality spinal cord fMRI data. First, in all three MRI studies, we made use of a custom-built cervical spinal cord coil with 24 receive channels (Figure 1A, 1B), the arrangement of which allowed for a high in-plane acceleration. We were able to record EPI data with a reduction factor of 3, leading to reduced distortions and signal loss (Figure 2A); note that this coil also performed well with regards to acceleration in recent 7T benchmarking coil tests (Alonso-Ortiz et al., 2025). We furthermore pursued a passive shimming approach, using a flexible and comfortable collar around the neck filled with susceptibility-matched material (Figure 1A). This resulted in a substantial reduction in large-scale magnetic field inhomogeneity in the spinal cord region: data from a single participant are depicted in Figure 2B, with group data (N = 7) also showing a consistent improvement in B_0_ field homogeneity when comparing acquisitions with and without passive shimming (Supplementary Material).

**Figure 1.**
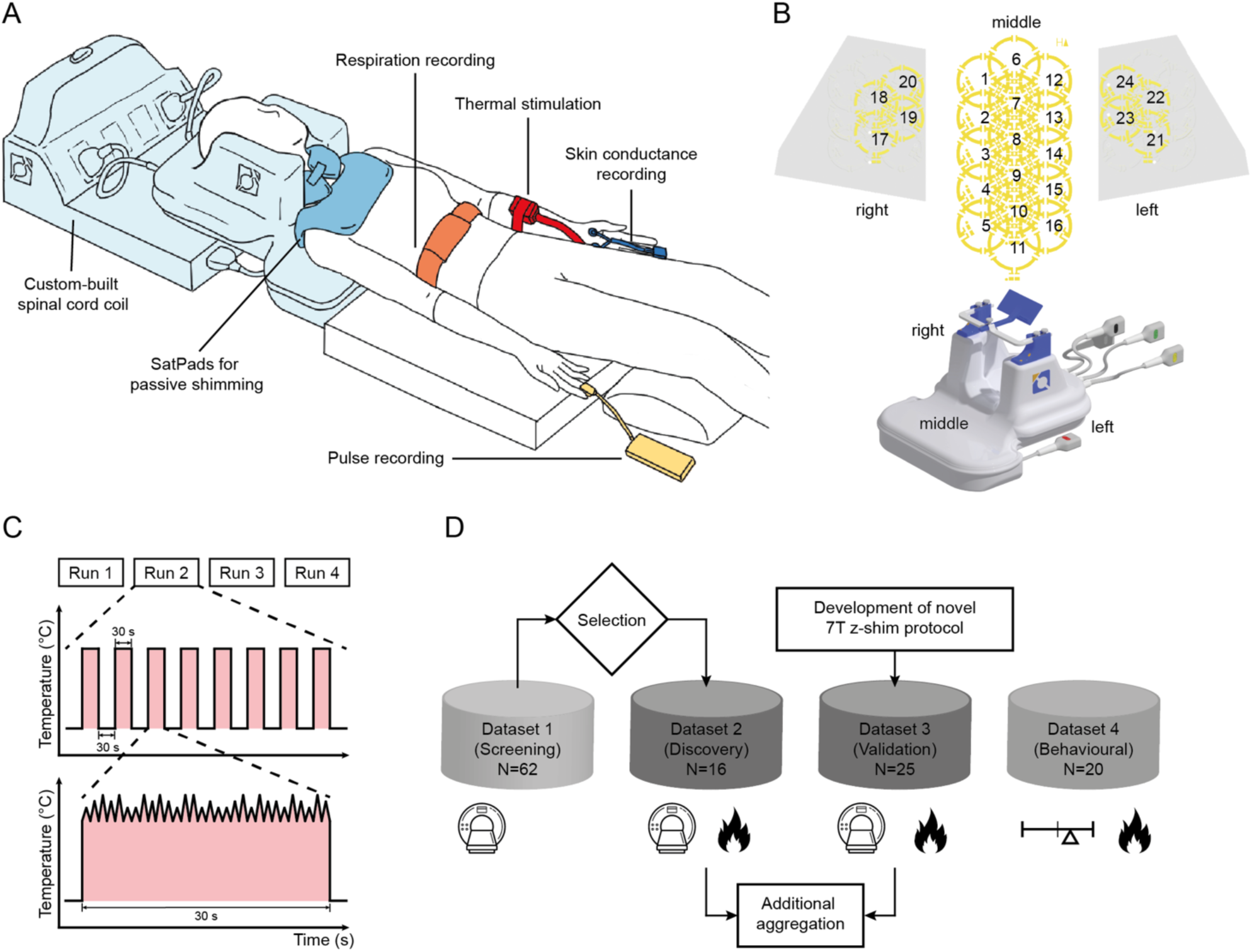
Experimental approach. **A.** Schematic overview of experimental setup for MRI, depicting all equipment used. Participants’ head and neck rested in a custom-built 24-channel neck coil for cervical spinal cord imaging and susceptibility matched SatPads for passive shimming were placed around their neck and over their chest. In Datasets 2 and 3, thermal stimulation occurred on the left forearm via an MR-compatible thermode and respiratory and cardiac data were recorded to correct fMRI data for physiological noise (skin conductance data were recorded as a physiological marker of pain in Dataset 2 only). **B.** The custom-built coil was anatomically formed and contained 24 receive channels: 3 rows along the neck (16 channels) and four channels on each side of the neck. **C.** The experimental heat-pain paradigm consisted of a robust block design (30 s ON, 30 s OFF) with four runs, each containing eight painful heat stimuli. Heat stimulation consisted of an individually determined plateau temperature (41°C - 45°C), superimposed with 30 pseudo-randomized heat spikes (2/3/4 °C), each < 1 s. **D.** Overview of datasets, including sample size and dependencies. Spinal cord MRI data were acquired in Datasets 1-3, fMRI data in Datasets 2-3, and the heat pain paradigm was used in Datasets 2-4. Dataset 1 is a *screening* dataset, from which participants with adequate data quality were selected for Dataset 2, which is a *discovery* dataset, allowing for exploring heat-pain induced spinal cord BOLD responses. Dataset 3 is a preregistered *validation* dataset, replicating the paradigm from Dataset 2 in an independent participant cohort with a novel 7T EPI sequence; both Datasets show similar results, and their results are shown in aggregated form as well as separately (Supplementary Material). Dataset 4 is a follow-up *behavioural* dataset (i.e. acquired without MRI), allowing for exploring supraspinal response components to the employed heat-pain stimulation.

**Figure 2.**
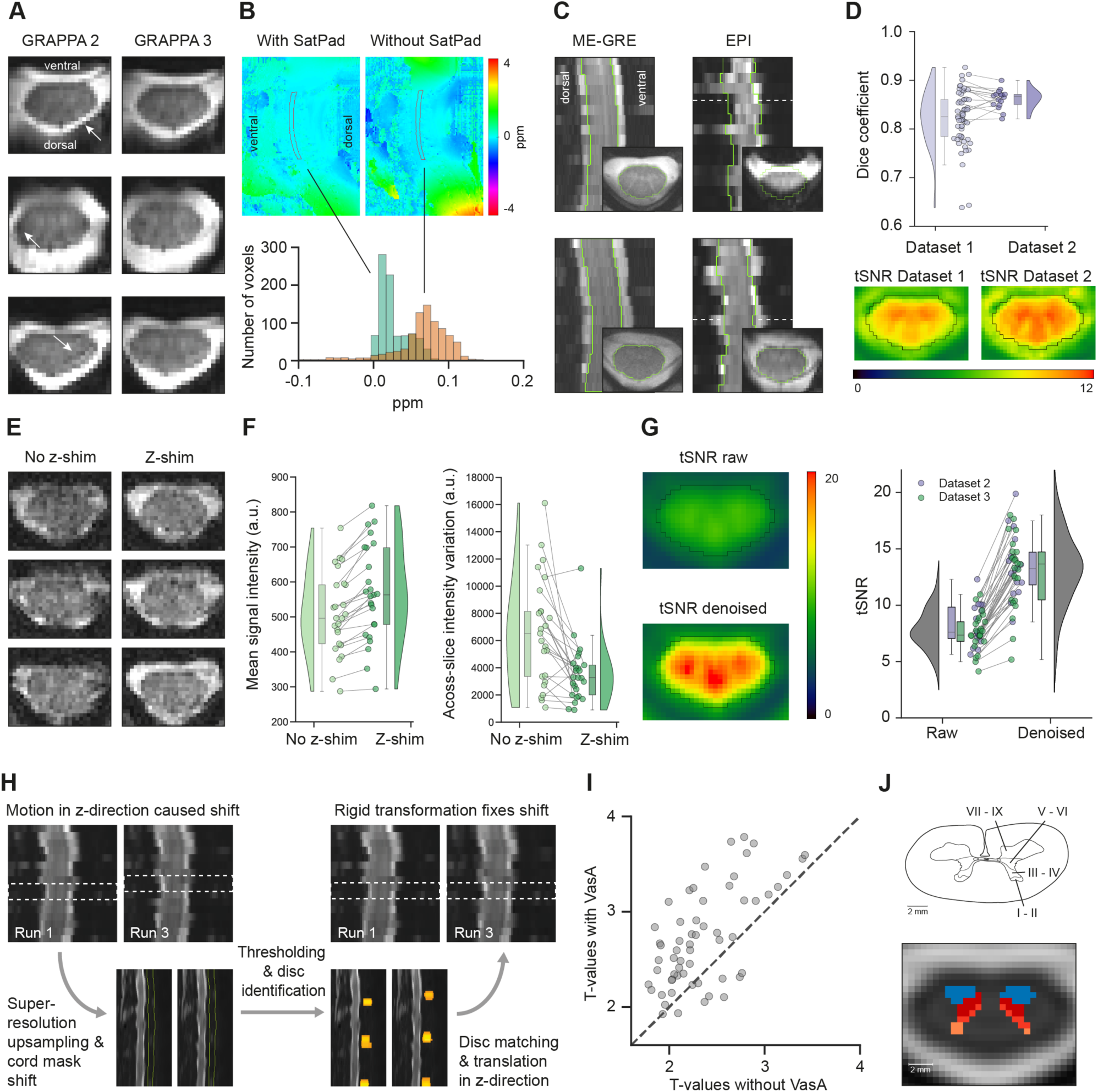
Methodological improvements. **A.** The coil design supports EPI data acquisition with an in-plane acceleration (via GRAPPA) of up to factor 3, with exemplary EPI slices demonstrating reduced distortions and signal loss (compared to factor 2; see also exemplary arrows). **B.** Data acquisition with passive shimming via a susceptibility matched collar reduces large scale field inhomogeneities, as demonstrated in the exemplary sagittal field-map section (top panel; spinal cord outlined in grey) and the extracted spinal cord data (bottom panel). **C.** The selection of participants from Dataset 1 for Dataset 2 was based on EPI data quality. Exemplary images from two participants (anatomical ME-GRE and functional EPI) highlight inter-individual differences in EPI data quality between selected participants (bottom) and those not selected (top) with signal loss and distortion observed in functional images). The green outline depicts the ME-GRE spinal cord segmentation and the dotted line denotes the transverse slice location. **D.** EPI data quality (exemplified visually in panel C) was evaluated using Dice coefficients (we assessed the overlap of cord segmentations derived from functional and anatomical images (top)) and tSNR maps of unprocessed functional images (bottom). Consistent data across sessions ensures the high quality of Dataset 2. **E.** Exemplary single-volume slices from Dataset 3 demonstrate that slice-specific z-shimming improved signal intensity (right) compared to no z-shimming (left). **F.** Slice-specific z-shimming increased the average spinal cord fMRI signal intensity (left) and reduced the signal intensity variation across slices (right); dots indicate participants. **G.** MP-PCA denoising was employed to reduce thermal noise in EPI data and thus enhance spinal cord tSNR in cord (depicted are a group-average C6 cross-section in template space, with cord location outlined (left) and individual data points across Datasets 2 & 3 (right). **H.** To improve motion correction, we developed a procedure for detecting and correcting motion in the through-slice direction, based on disc alignment. **I.** Vascular auto-rescaling (VasA) leads to a sensitivity increase in group-level statistics (depicted are t-values from group-level analyses in the target region C6 left dorsal horn with and without VasA). **J.** Based on an annotated C6 spinal cord cross section (Sengul et al., 2013), we created a high-resolution layer-specific grey matter atlas in PAM50 space to allow for layer-specific examination of BOLD responses.

The quality of single-shot gradient-echo EPI data in the spinal cord varies strongly across participants at ultra-high field strength, due to inter-individual differences in local magnetic field inhomogeneities, for example leading to strong signal loss in some participants. Since this prevents high-fidelity fMRI at the group level, we addressed this problem via two different strategies. In a first approach, we acquired a large screening dataset (Dataset 1, N = 62), from which we selected participants with adequate data quality (i.e. low levels of distortions and signal loss in EPI data) for participation in the task-based fMRI study (Dataset 2, N = 16). This selection was possible because data-quality differences across participants (examples in Figure 2C: upper row with ‘poor’ data, lower row with ‘good’ data) are stable across time. The selected participants showed substantially lower levels of distortions and signal loss (as assessed by Dice coefficients between anatomical multi-echo gradient-recalled echo (ME-GRE) and functional EPI data; Figure 2D, upper panel; average dice coefficient in Dataset 1: 0.82 and in Dataset 2: 0.86) and higher temporal signal-to-noise ratio (tSNR; Figure 2D, lower panel; average tSNR in cord in Dataset 1: 8.30 and in Dataset 2: 9.01). While this approach allows for obtaining high quality fMRI data, it is highly time-consuming.

Therefore, in a second approach we developed an EPI protocol for automated slice-specific z-shimming, which allowed for the acquisition of an independent high-quality dataset without the need to pre-select participants (Dataset 3; N = 25). This technique is based on earlier work at 3T (Finsterbusch et al., 2012; Kaptan et al., 2022) and was now optimized for 7T, e.g. by adding a GRE-based autocalibration method to allow for robust parallel imaging (Seifert & Xu, 2022). With slice-specific z-shimming, we observed substantial reductions in signal loss and signal variability across slices, as demonstrated visually in single EPI-volumes (Figure 2E) and statistically in group-level data (Figure 2F; signal intensity: t(24) = 8.89, p = 4.7·10^-9^, d = 1.78; signal intensity variation: t(24) = 4.05, p = 0.0005, d = 0.81). Taken together, the use of a novel coil design in combination with passive shimming and participant pre-selection or slice-specific z-shimming provided the basis for obtaining high-quality spinal cord fMRI data.

We then aimed to further refine these data with tailored improvements in data analysis, where we combined several approaches for increasing the sensitivity and spatial precision of our data. First, considering that increased in-plane acceleration and small voxel sizes as used here lead to thermal noise dominated fMRI data (Vizioli et al., 2021), we employed a thermal noise correction approach using Marchenko-Pastur Principal Component Analysis (MP-PCA). This resulted in a substantial increase in tSNR (Figure 2G; Dataset 2: tSNR increase of 63.4%, t(15) = 15.15, p = 1.69·10^-10^, d = 3.79; Dataset 3: tSNR increase of 71.0%, t(24) = 13.72, p = 7.39·10^-13^, d = 2.74), but did not lead to a substantial penalty in terms of spatial smoothness (Dataset 2: increase in full width at half maximum (FWHM) from 0.92 mm to 1.02 mm; Dataset 3: increase in FWHM from 1.00 mm to 1.13 mm). Second, we leveraged our fMRI data acquisition’s higher resolution in z-direction (compared to typical 3T studies) and developed a new registration procedure for EPI data to correct movements in the foot-head direction between runs by registering the intervertebral discs onto another (Figure 2H). Such movements would remain uncorrected with typical in-plane motion correction procedures. Third, we accounted for vascular reactivity differences between participants by using a vascular autorescaling technique (VasA; Kazan et al., 2016, 2017) that improved statistical sensitivity at the group level (16.6% increase in t-scores in the left dorsal horn of segment C6; Figure 2I). Finally, we aimed to capitalize on the higher resolution offered by 7T fMRI (approximately 0.8 × 0.8 mm^2^ in-plane resolution in our work) and therefore created a layer-specific grey matter spinal cord MRI atlas (Figure 2J). This was based on post-mortem histological data available via the human part of the Atlas of the Spinal Cord (Sengul et al., 2013) and allowed us to delineate the layer-specific distribution of BOLD responses in the dorsal horn. Collectively, these improvements allowed us to interrogate spinal cord fMRI data at high resolution with high sensitivity.

### 2.2. Assessing group-level spinal cord responses to 30 s heat stimuli

In a first set of group-level analyses, we tested whether we would be able to detect significant spinal cord BOLD responses in response to 30 s heat stimuli in an fMRI block design. Following a pre-registered analysis plan, we report results for the combined dataset (pooling Datasets 2 and 3; main manuscript), as well as for each dataset separately (Supplementary Material). At an exploratory threshold (p < 0.01 uncorrected), we observed widespread activation that i) spanned several spinal segments rostro-caudally with multiple peaks, although the largest peak occurred in target segment C6 (Figure 3A) and ii) was found primarily on the ipsilateral (left) side but clearly extended contralaterally. When using strict statistical thresholding (family-wise-error correction at p<0.05), two clusters were observed in the target region of the left dorsal horn in segment C6 (cluster 1: 35 voxels, peak voxel: t = 3.78, p = 0.004; cluster 2: 11 voxels, peak voxel: t = 3.56, p = 0.012; Figure 3B; for separate results in Datasets 2 and 3, see Figure S1).

**Figure 3.**
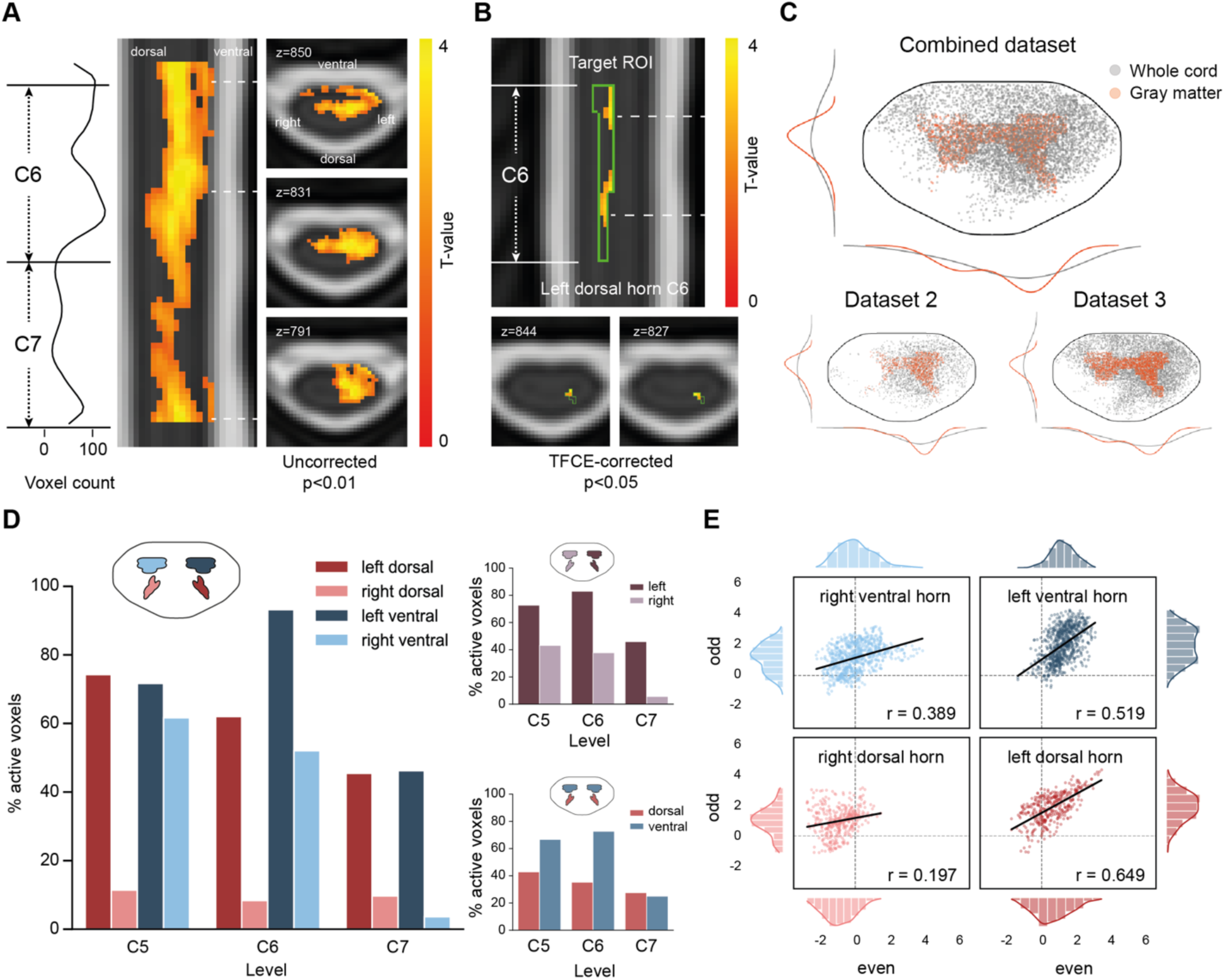
Group-level spinal cord BOLD responses. **A.** Uncorrected group-level results (p<0.01) constrained within the spinal cord for the combined dataset (Datasets 2 & 3) in response to 30 s heat pain. The density plot (left) shows the rostro-caudal activation distribution, with segments C6 and C7 marked. The sagittal view (middle) shows activation spread across spinal levels with the dashed lines indicating the position of the axial plots (right) at maximum activation in cord segments C5, C6, and C7. **B.** TCFE-corrected results (p_FWE_<0.05) for the combined dataset in response to 30 s heat pain in the target ROI (C6 left dorsal horn, green outline) show two clusters with peaks depicted in axial plots. **C.** Unconstrained depiction of the spatial distribution of BOLD responses: single-slice projected positions of suprathreshold voxels (uncorrected p<0.05) across segments C5 - C7 in whole spinal cord (grey dots) and grey matter only (red dots), with jitter added for visualization. The outside density plots show the voxel distributions across hemicords. Results are depicted for the combined dataset (top) and for Datasets 2 & 3 separately (bottom). **D.** Percentage of suprathreshold voxels (uncorrected p<0.05) in each grey matter horn mask (left) and separated into either left and right masks (right top) or dorsal and ventral masks (right bottom). **E.** Split-half reliability based on the t-values of all voxels within a grey matter horn mask, estimated via separate GLMs for even and odd trials (depicted are all four horns, with dashed lines indicating zero and histograms depicting the t-value distributions).

Next, we aimed to provide a more holistic picture of spinal cord activation. First, we projected all activated voxels onto one transverse spinal cord plane (Figure 3C; p < 0.05 uncorrected) and – via this unconstrained depiction – observed i) responses across both the white matter (grey dots) and grey matter (red dots), ii) a bilateral activation with ipsilateral predominance, iii) the highest activation density in the central part of the dorsal-ventral axis. Activation patterns are consistent across datasets, with Dataset 3 showing more contralateral extension, possibly due to higher power (N = 25 vs. N = 16), yet a left hemi-cord preference remains. While widespread, the activation patterns (at p < 0.05 uncorrected) seem primarily concentrated in grey matter: 56.4% of grey matter voxels are activated (average t-value of 2.36) and 26.4% of white matter voxels are activated (average t-value of 1.96). Second, we provide count-based summary measures (Figure 3D) of suprathreshold voxels within each horn and – via this grey-matter constrained analysis – observed that i) all three segments showed clear responses, ii) there was a strong ipsilateral predominance, and iii) more extensive ventral than dorsal activation was evident.

Finally, we complemented the above analyses focused on activation strength and spatial distribution by an approach that addresses response stability (i.e., reliability). We performed an even-odd split of the data, estimated separate general linear models (GLMs) for even and odd trials, and then assessed reliability by correlating the group-level t-statistics of all voxels within each of the four grey matter regions for even and odd data. The most reliable spatial pattern was observed in the ipsilateral dorsal horn, with a large effect size (r = 0.65), higher than in the contralateral dorsal horn (r = 0.20) and the ventral horns (r = 0.39 and r = 0.52). Overall, these analyses thus provide an overarching characterization of BOLD responses across the spinal cord and demonstrate their significance and reliability in the ipsilateral dorsal horn, i.e. the site of the first synaptic relay in the nociceptive system.

### 2.3. Investigating individual activation patterns

Next, we capitalized on the higher SNR available at 7T and aimed to investigate the individual activation patterns underlying the above-reported group result. The group-level results are confined to spinal segments C6 to C7 (with a slight contribution by segment C5), where all participants have data in template space, though individual coverage may vary due to anatomy (Figure 4A). We observed that individual activation maxima in the left dorsal horn exhibited a high variability in their rostro-caudal position, spanning segments C5 to C8 (Figure 4B). Since a single peak voxel might be insufficient to gain insight into activation profiles, we also present each participant’s complete rostro-caudal activation pattern in the left dorsal horn (Figure 4C). This depiction shows widespread activation across segments, with high inter-individual differences and often multiple peaks per participant in the rostro-caudal direction. Such inter-individual variability allows for two interpretations: either these highly variable patterns occur due to noise or they reflect stable idiosyncratic responses.

**Figure 4.**
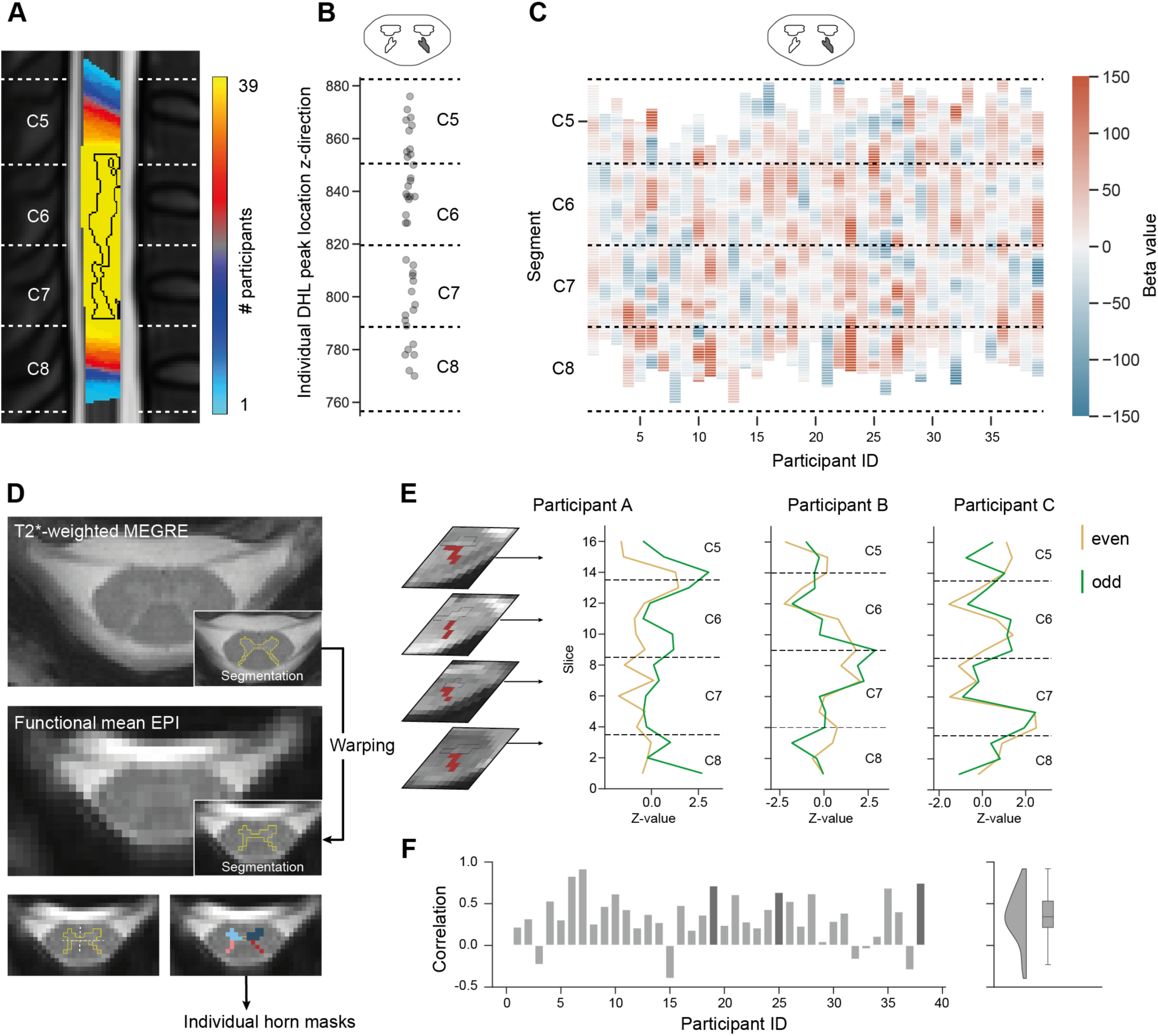
Individual-level spinal cord BOLD responses. **A.** Depicted is the uncorrected group-level result from Figure 3 (via outline in black) overlaid on a sagittal slice where colours indicate overlap of data acquisition coverage across participants; individual datasets vary in segmental coverage (white lines indicate segmental [and not vertebral] borders). **B.** Individual peak activations (maximal parameter estimate, i.e. beta value) in the left dorsal horn occur at different spinal segments. **C.** Individual activation patterns (average left dorsal horn beta estimate per slice in template space) show inter-individual response spread across spinal segments. **D.** Depicted is one participant’s high-resolution T2*-weighted ME-GRE image (top panel) with segmented grey matter (inset). This is then warped into EPI space (middle panel), where an automated determination of midline and dorsal/ventral horn borders enables participant-specific creation of grey matter horn masks (bottom panel). **E.** Results from exemplary participants, with lines showing slice-wise BOLD signals (average z-values) from individual left dorsal horn masks for even (yellow) and odd (green) trials; for one participant, the individual dorsal horn mask is also shown in exemplary slices. Note that participants display slightly different segmental coverage and clearly different peak location but reliable activation, as evidenced by the similarity of the even and odd activation profile across slices. **F.** Correlation values, as computed from even and odd slice-wise dorsal horn BOLD signals (average z-value per slice), for all individual participants (left panel, participants displayed in E are highlighted with dark grey bars) and summarized across the group (right panel). Note that the analysis in this panel is based on 38 participants, as one participant lacked a T2*-weighted ME-GRE image.

We therefore assessed individual participant’s responses in more detail in their native space. To precisely segment each individual’s spinal cord grey matter, we employed the high-resolution T2*-weighted ME-GRE images, which were reconstructed offline using a recently developed optimized navigator-based correction approach for increased SNR (Beghini et al., 2025). The obtained individual grey matter segmentation was registered to the EPI space, where an automated procedure detected the midline and dorsal/ventral horn borders, allowing for the creation of individual horn masks (Figure 4D). We then extracted the average activation values from the left dorsal horn across all slices for even and odd trials and, for each individual, estimated the across-slice activation pattern correlation. Exemplary participants display activation peaks at different spinal segments, but in a reliable way across even and odd trials (Figure 4E). While this slice-profile correlation naturally varies across participants, its group-level average has at least a medium effect size (r = 0.338; Figure 4F) with individual datasets showing similar effect sizes (Dataset 2: r = 0.386; Dataset 3: r = 0.305). In summary, these analyses demonstrate substantial inter-individual spatial variability (which cannot be inferred from group results alone), that is however reliable within participants and thus of neurobiological interest.

### 2.4. Probing for multiple response types

So far, all analyses were confined to interrogating BOLD responses that follow the ON-OFF 30s block design structure of our heat-pain paradigm. However, in non-human primates, such prolonged heat stimulation typically results in two different types of spiking in nociceptive Aδ nerve fibres in the peripheral nervous system (schematically depicted in Figure 5A; Treede et al., 1995, 1998): a sustained firing of action potentials throughout the stimulus (‘Type I heat response’) and a phasic firing at onset that quickly diminishes after ∼3 s (‘Type II heat response’); note that a similar distinction is observed for C-fibres (Johanek et al., 2008; Meyer & Campbell, 1981). We therefore probed for a second response type in addition to the ‘sustained response’ described above (modelled using the entire 30 s, trying to match Type I), by testing for a ‘phasic response’ at stimulus onset (modelled using the first 3 s, trying to match Type II; Figure 5B).

**Figure 5.**
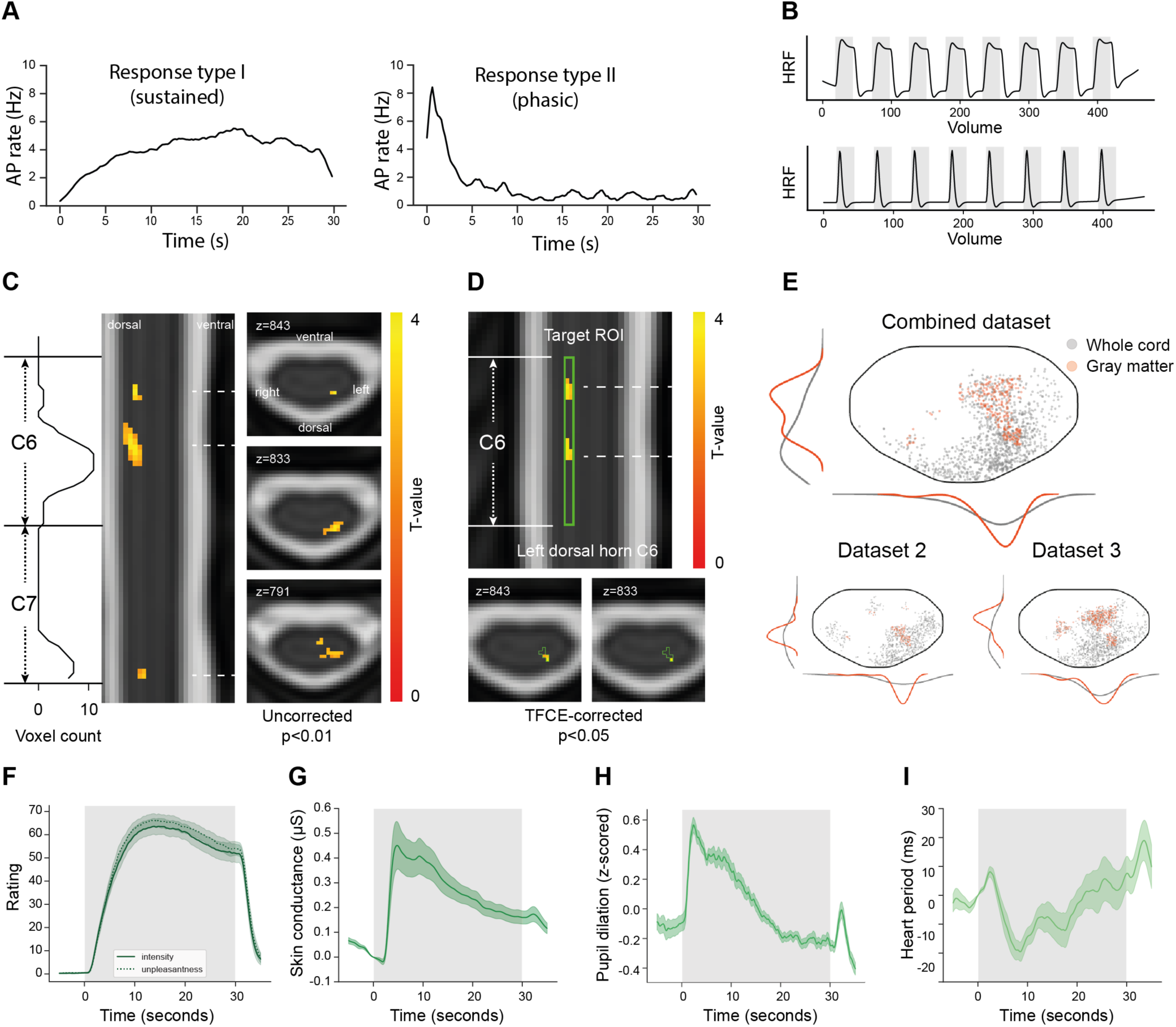
Different response types to heat pain stimulation. **A.** Two peripheral nerve response types from previous work in monkeys (simplified depiction from Treede et al., 1998): action potential rates to 30 s heat stimulus show sustained (left) and phasic responses (right) in different Aδ fibres. **B.** Spinal cord fMRI heat response GLMs: the sustained response is modelled using a 30 s regressor (for results see Figure 3) and the phasic response is modelled using a 3 s regressor per pain stimulus. **C.** Uncorrected group-level results (p<0.01) in the spinal cord for the combined dataset (Datasets 2 & 3) using the phasic response GLM. The density plot (left) shows the rostro-caudal activation distribution, with segments C6 and C7 marked. The sagittal view (middle) shows activation spread across spinal segments with the dashed lines indicating the position of the axial plots (right) at each cluster’s peak. **D.** TCFE-corrected results (p_FWE_<0.05) for the combined dataset’s phasic BOLD response in the target ROI (C6 left dorsal horn, green outline) show two clusters with peaks depicted in axial plots. **E.** Unconstrained depiction of the spatial distribution of phasic BOLD responses: single-slice projected positions of suprathreshold voxels (uncorrected p<0.05) across segments C5 -C7 in whole spinal cord (grey dots) and grey matter only (red dots), with jitter added for visualization. The outside density plots show the voxel distributions across hemicords. Results are depicted for the combined dataset (top) and for Datasets 2 & 3 separately (bottom). **F.-G.** Data from the behavioural experiment (Dataset 4, N = 20), showing group-average **F.** continuous pain rating (intensity = solid line, unpleasantness = dotted line), **G.** skin conductance response, **H.** pupil dilation response, and **I.** heart period response over the duration of the heat stimulus (shaded grey area).

We indeed observed a second, phasic, response type in the spinal cord. Similar to the sustained response, a large part of the activation was present in target segment C6, though in a sparser way in terms of rostro-caudal extent and also with a more dorsally centred pattern (Figure 5C, p < 0.01 uncorrected); similar patterns were observed in both datasets (Figure S2). When using strict statistical thresholding (family-wise-error correction at p < 0.05), two clusters were observed in the target region of the left dorsal horn in segment C6 (cluster 1: 9 voxels, peak voxel: t = 4.03, p = 0.0068; cluster 2: 6 voxels, peak voxel: t = 4.03, p = 0.0068; Figure 5D). When assessing the entire activation pattern in an unconstrained way (projection onto one transverse spinal cord plane), we observed the phasic response to be more lateralized than the sustained response and to also have a stronger focus towards the tip of the dorsal horn (Figure 5E); this replicated across both datasets.

Having uncovered this novel evidence for two different response types at an early stage of human central nervous system processing, we next aimed to assess whether such response components would also be evident in higher-level subjective and autonomic metrics of pain processing, as this might give hints towards the behavioural relevance of these spinal responses. We therefore carried out a separate behavioural experiment outside the MRI (Figure 1D, Dataset 4, N = 20) with the same pain paradigm. Participants continuously rated intensity and unpleasantness of heat stimulation in separate runs and had their skin conductance, pupil dilation and heart-period responses recorded. Both types of ratings showed a sustained response profile (Figure 5F; intensity: p = 0.0003, significant cluster: 0 s – 30 s; unpleasantness: p = 0.0001, significant cluster: 0 s – 30 s). While skin conductance responses peaked early and then remained sustained above baseline for the whole stimulus period (Figure 5G; p = 0.0001, significant cluster: 2.6 s – 30.0 s; Figure S3A), pupil dilation responses peaked very early and diminished quickly, thus showing a phasic response (Figure 5H; p = 0.0001, significant cluster: 1.0 s – 13.9 s). Heart period responses were harder to assign to a sustained or phasic response mode, since they followed a triphasic pattern in the first half of the stimulus (Figure 5I; Figure S3B). Collectively, we have thus established evidence for multi-faceted spinal cord BOLD responses to prolonged heat-pain stimulation, which is paralleled by higher-level subjective and autonomic responses.

### 2.5. Delineating layer-specific spinal cord responses

Finally, we leveraged the higher resolution offered by our ultra-high-field imaging approach (with an approximately three-fold voxel volume reduction compared to typical 3T spinal cord fMRI voxels) to investigate BOLD responses in different spinal cord layers, which are the fundamental processing units within the dorsal horn. To achieve this, we made use of an annotated post-mortem cross-section of a human spinal cord, where histological analyses allowed for precise identification of the laminar structure of the spinal cord (Sengul et al., 2013). The annotated image was digitized, rasterized, transformed to a NIfTI file and then warped to the coordinate space of the Spinal Cord Toolbox, where it was employed for distinguishing superficial (laminae I & II), middle (laminae III & IV), and deep layers of the dorsal horn (laminae V & VI), as well as the ventral horn (laminae VII – IX) at the level of segment C6 (Figure 6A). While both superficial and deep dorsal horn layers have traditionally been associated with heat pain responses, here we explored whether the sustained and phasic BOLD response might differentially map onto layers.

**Figure 6.**
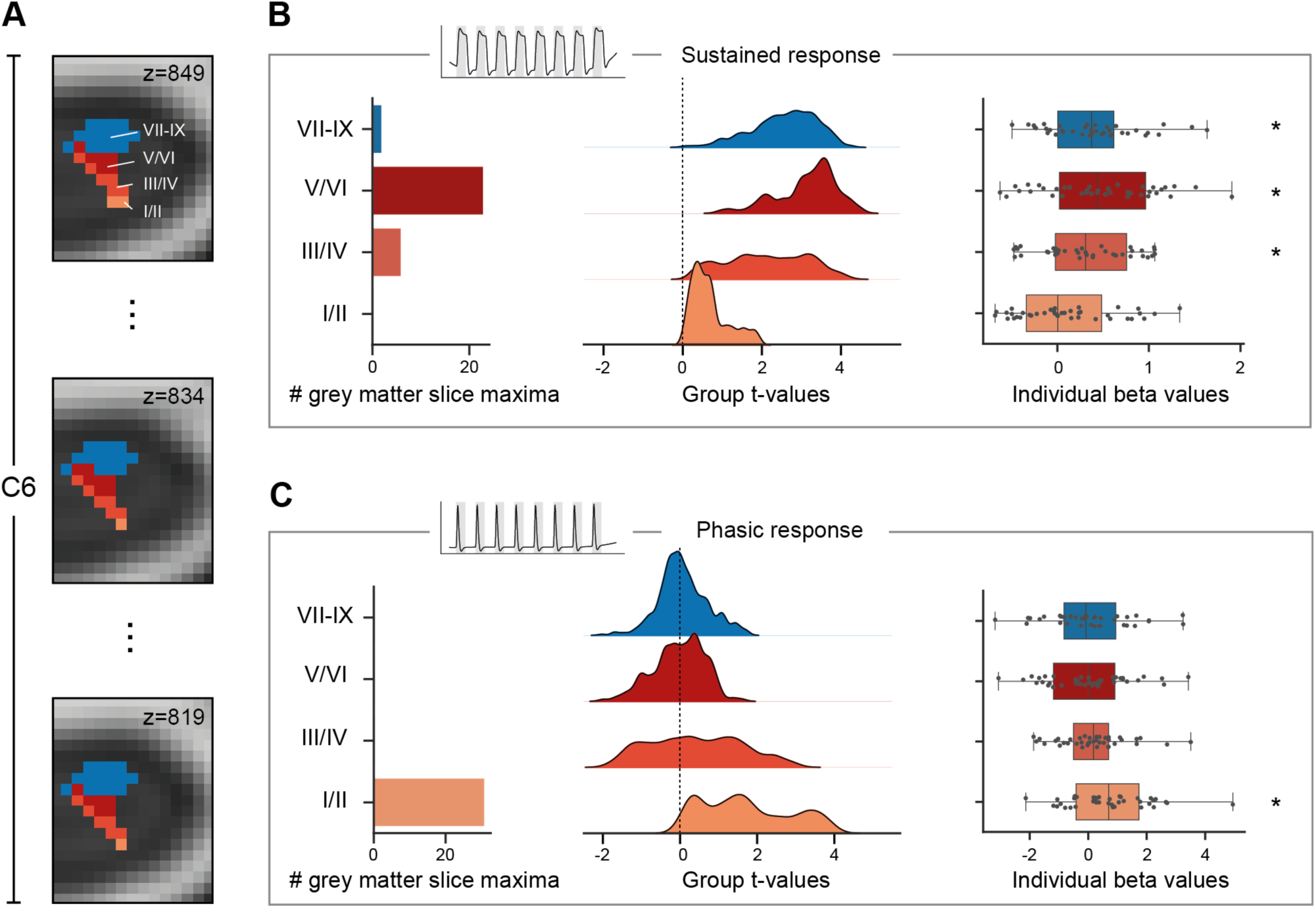
Layer-specific differentiation of spinal cord BOLD responses. **A.** A newly-created and histology-based atlas in PAM50 template space distinguishes layers within the grey matter (depicted is the left hemicord): superficial dorsal horn (laminae I/II), middle dorsal horn (laminae III/IV), deep dorsal horn (laminae V/VI), and ventral horn (laminae VII-IX); three (out of 31) exemplary C6 slices are displayed. **B.** Layer-specific BOLD responses for the sustained heat GLM: slice-wise occurrence of the group result’s maximum grey matter voxel in each layer (left); distribution of group analysis t-values in each layer (middle); individual participants’ average activation (beta) values in each layer (right; asterisk marks significance for test against 0). **C.** Layer-specific BOLD responses for the phasic heat GLM, with panels organized as in B. All results in B and C are based are based on minimally smoothed data, with unsmoothed data showing virtually identical results (Figure S4).

Analysing the sustained BOLD response with this atlas revealed that the slice-wise group-level activation peaks most frequently occurred in the deep layers of the dorsal horn compared to all other grey matter regions (Figure 6B, left panel); especially noteworthy is the absence of peaks in the superficial layers. When examining voxel-wise group-level t-statistics across all C6 slices, the t-value distribution in the deep dorsal horn layer was shifted to the right and strongly right-skewed (indicating a predominance of high t-values), while the distribution of t-values in the superficial layers was centred close to zero (Figure 6B, middle panel). At the participant level, average individual beta values per layer revealed that only the superficial dorsal horn layer activation did not significantly differ from zero (t(38) = 1.22, p = 0.16, d = 0.20), with the deep dorsal and ventral horn showing the most prominent effects (Figure 6C, right panel; middle dorsal horn: t(38) = 4.11, p = 0.0001, d = 0.66; deep dorsal horn: t(38) = 4.84, p = 1.10·10^-5^, d = 0.77; ventral horn: t(38) = 4.88, p = 9.62·10^-6^, d = 0.78; all tests one-tailed). To contrast the activation in superficial and deep dorsal horn, we compared the individual beta values against each other, observing significantly weaker activation in the superficial compared to the deep dorsal horn (t(38) = -3.31, p = 0.002, d = -0.53; two-tailed).

In contrast, analyses of the phasic BOLD response demonstrated that the group-level activation peak per slice occurred exclusively in the superficial layers (Figure 6C, left panel). Here, all t-value distributions were centred around zero, except for those in the superficial layers (Figure 6C, middle panel). At the individual level, only activation in the superficial dorsal horn significantly differed from zero (Figure 6C, right panel; superficial dorsal horn: t(38) = 3.19, p = 0.001, d = 0.51; middle dorsal horn: t(38) = 0.97, p = 0.17, d = 0.16; deep dorsal horn: t(38) = -0.04, p = 0.51, d = -0.006; ventral horn: t(38) = 0.15, p = 0.44, d = 0.02; all tests one-tailed). The superficial dorsal horn activation was significantly stronger than the deep dorsal horn activation (t(38) = 2.59, p = 0.013, d = 0.42; two-tailed), highlighting the profoundly different activation patterns for the different response types. These results replicated to a very large degree across several control analyses, i.e. i) analysing Dataset 2 and Dataset 3 separately, ii) investigating unsmoothed data, and iii) carrying out non-parametric permutation testing on voxel-wise data (see Supplementary Material).

Taken together, these high-resolution fMRI data provide several lines of evidence for fine-grained processing of nociceptive inputs at the first central nervous system stage. Phasic responses occur in superficial (but not deep) layers and sustained responses occur in deep (but not superficial) layers of the dorsal horn, indicating a layer-specific double-dissociation.

## 3. Discussion

In this study, we developed the methods for and demonstrated the feasibility of layer-specific UHF fMRI in the human spinal cord. Our method thus provides mesoscopic access to sensory information processing at the first synaptic relay of the CNS in humans. Using a data acquisition strategy encompassing screening, discovery and preregistered validation datasets, we were able to obtain robust and replicable insights into the processing of nociceptive stimuli in the dorsal horn. Notably, we discovered i) reliable individual response patterns, ii) temporally distinct nociceptive responses to the same input, and iii) a precise layer-specific organization of spinal BOLD responses.

While MRI data acquisition and analysis methods have become increasingly standardized for spinal cord measurements at 3T (Cohen-Adad et al., 2021; De Leener et al., 2017, 2018), technical challenges have hindered progress in spinal cord fMRI at higher field strengths and thus prevented mesoscopic measurements of human spinal cord function (Barry et al., 2017). Due to this limitation, high-resolution insights into spinal processing remain the exclusive domain of research in animal models, a longstanding barrier we aimed to overcome here via numerous methodological improvements. The use of a custom-built coil and a passive shimming approach reduced distortions of EPI data and increased B_0_ homogeneity, respectively. Yet, despite these efforts, the screening dataset demonstrated EPI data quality still varying substantially across individuals, preventing UHF fMRI studies with a standard sample of participants. One successful approach to address this variability (taken in the discovery dataset), was to rigorously pre-select participants based on their EPI data quality in the screening dataset, but this is resource intensive and may bias population sampling. For the validation dataset, we thus pursued a time-efficient and unbiased approach by applying an UHF-optimized implementation of automated slice-specific z-shimming (Finsterbusch et al., 2012; Kaptan et al., 2022), leading to large effect sizes in terms of signal recovery and reduction of signal variation across slices.

These novel UHF data acquisition strategies resulted in two high quality datasets, which we then refined using advanced denoising techniques in order to further push the boundaries of current imaging capabilities. First, we employed a thermal noise correction method that increased tSNR by over 60% without compromising spatial resolution. Second, physiological noise, particularly prominent in the spinal cord, was extensively corrected using multiple approaches. Third, we optimized group-level statistical sensitivity by addressing participant variability in vascular reactivity through a validated rescaling strategy. Ultimately, we then leveraged these advances to assess BOLD responses in the dorsal horn, the small size of which typically leads to strong partial volume effects, making it optimally suited for high-resolution imaging at 7T.

Using a heat-pain block design, we identified a robust activation pattern within the ipsilateral dorsal horn that remained significant even after rigorous statistical correction. While such robust heat-pain responses have not previously been observed at 7T, they align well with previous 3T studies (Brooks et al., 2012; Dabbagh et al., 2024; Eippert et al., 2009; Geuter & Büchel, 2013; Nash et al., 2013; Oliva et al., 2022; Sprenger et al., 2012, 2015, 2018; Summers et al., 2010; Tinnermann et al., 2017; Weber et al., 2016a; see Kolesar et al., 2015 for a review and Seifert et al., 2024 for early 7T work). We observed extensive activation across the cross-section of the cord: while there was a pronounced ipsilateral dominance, consistent with previous studies, contralateral activity was also observed (Cahill & Stroman, 2011; Dabbagh et al., 2024; Geuter & Büchel, 2013; Seifert et al., 2024; Sprenger et al., 2018; Summers et al., 2010; Weber et al., 2016a; Yang et al., 2015). The most pronounced activation was observed in our target segment C6, but activation was also present in neighbouring segments, fitting with both previous fMRI studies (Dabbagh et al., 2024; Rempe et al., 2015; Seifert et al., 2024; Sprenger et al., 2018; Weber et al., 2016a), and invasive recordings in animal models (Kato et al., 2004; Shekhtmeyster, Duarte, et al., 2023). Most importantly, when performing a split-half analysis, we observed the highest reliability of BOLD response patterns (with a large effect size) in the ipsilateral dorsal horn, i.e. the primary site of synaptic nociceptive processing, speaking for the robustness of our novel imaging approach.

Despite this high reliability, some sources of interindividual variability cannot be eliminated and might be of neurobiological importance. We therefore made use of the increased SNR at 7T as well as the high-resolution ME-GRE data we acquired (allowing for precise grey matter segmentation in individuals) and investigated the individual participant dorsal horn activation patterns underlying the group results. We discovered that rostro-caudal activation location can differ substantially between individuals and that these idiosyncratic differences in BOLD activation profiles are reliable, thus speaking against them arising due to noise and fitting with similar variations detected in optical recordings in animal models (Sasaki et al., 2002). Variability in dermatomal location (M. W. L. Lee et al., 2008) or nerve root termination (Cadotte et al., 2015) might be factors contributing to these differences and developments in nerve-root identification could shed light on this question (Valošek et al., 2024). Given that our results indicate the feasibility of individual-participant spinal cord fMRI at 7T, future studies based on dense-sampling approaches could investigate pain variability across time in the healthy state or monitor disease progress / treatment response in individuals with chronic pain or other neurological disorders with spinal pathology (e.g. multiple sclerosis or spinal cord injury).

We also investigated whether the spinal cord’s response might be more complex than demonstrated by previous fMRI studies and indeed found initial evidence for multiple spinal responses when also detecting a phasic BOLD response occurring at the onset of stimulation. Such a temporal dissociation has previously been observed in electrophysiological peripheral nerve recordings of nociceptive primary afferents in non-human primates (Johanek et al., 2008; Treede et al., 1998) and in calcium imaging of the dorsal root ganglion in rodents (Qi et al., 2024; F. Wang et al., 2018). However, the existence of such response differentiation in the spinal cord, i.e. the projection target of these primary afferents, was hitherto unknown. To assess the functional relevance of such two-fold responses, we explored whether supraspinal indices of pain processing would also exhibit sustained and phasic response patterns. This was the case: while unpleasantness and intensity ratings showed sustained responses, pupil dilation exhibited a phasic response pattern (with skin conductance displaying a mixture of both patterns). These consistent spinal and supraspinal responses underscore the need to assess the temporal dynamics of nociceptive responses at multiple levels, as they likely play a crucial role in capturing the comprehensive experience of pain (Geuter et al., 2014). In a last step, we therefore aimed to leverage the unique capabilities of UHF fMRI (e.g. higher spatial resolution and shift in sensitivity towards smaller vessels; Uğurbil, 2014, 2018; Uludağ et al., 2009; Yacoub et al., 2001) to investigate the spinal expression of sustained and phasic responses at the mesoscopic scale.

By combining submillimetre fMRI data acquisition with the development of a novel fine-grained grey matter atlas, we were able to go beyond spinal cord fMRI studies at 3T – differentiating only dorsal and ventral horns – and investigated nociceptive processing across distinct functional units. More specifically, and in line with the concept that the dorsal horn should not be viewed as a single functional unit (Brown, 1982), we probed responses in spinal cord layers, which are a fundamental organizing principle across species (Rexed, 1952, 1954; Schoenen & Faull, 2004; Tan et al., 2023). By creating a layer-specific atlas in template space (though based on post-mortem data from a single individual), we made sure that our analysis strategy would be automated and easily reproducible, an important aspect for future routine use (considering that many layer fMRI approaches in the brain demand complex analysis tools and manual corrections; Chaimow et al., 2025). To address the potential drawback of template space group results not accurately representing individual activation patterns, we additionally provide individual participant data confirming the layer-specific group-level response pattern.

When analysing the layer-profile of the sustained BOLD response, we observed that the majority of grey matter activation is concentrated in the deep layers (V/VI) of the dorsal horn. Widespread activation in deep dorsal laminae in response to noxious stimulation has been widely reported in studies in rats, using extracellular recordings (Menétrey et al., 1977), c-Fos expression (Hunt et al., 1987; Presley et al., 1990; Williams et al., 1990) and autoradiographic methods (Abram & Kostreva, 1986; Coghill et al., 1991, 1993; Porro et al., 1991). In contrast, the phasic BOLD response at the onset of the heat stimulus almost exclusively targeted the superficial layers (I/II). Although the superficial layers are closer to the draining veins in the venous plexus (though separated by white matter; Thron, 2016), this response is unlikely to be an artifact because of the early nature of the response (with draining vein effects typically occurring in a delayed fashion; Kay et al., 2020), and because several control analyses confirmed the robustness of the layer-specific response dissociation. While the role of the superficial dorsal horn in nociceptive processing has been long established (Christensen & Perl, 1970; Kumazawa et al., 1975; Kumazawa & Perl, 1978; Light et al., 1979; Todd, 2010), the layer-specific response differentiation observed here is novel to our knowledge. It is supported however by in-vitro work of laminar-specific discharge rates (Ruscheweyh & Sandkühler, 2002) and autoradiographic and c-Fos studies in rats showing an early-to-late superficial-to-deep laminar shift (Hunt et al., 1987; Porro et al., 1991; Porro & Cavazzuti, 1992; Presley et al., 1990; Williams et al., 1990), though all of these used different time scales compared to our paradigm.

One possible explanation for the here-observed layer-specific response dissociation builds on “nociceptive specific” neurons being more prevalent in superficial layers and exhibiting relatively short-lived responses and “wide-dynamic-range” neurons being more prevalent in deep layers and exhibiting more tonic responses (Dougherty & Chen, 2016; Light, 1992; Lopez-Garcia & King, 1994), though obviously in different species and time scales. Linking to previous ideas (Ruscheweyh & Sandkühler, 2002; Sikandar et al., 2013), we speculate that the superficial layer responses observed here might thus link to novelty detection and induction of autonomic responses (e.g. pupil responses), whereas the deep layer responses might have a stronger link to the experience of sustained pain (e.g. subjective ratings). While we do not intend to draw oversimplistic conclusions with regards to the complexity of spinal processing (Light, 1992), it is a tantalizing prospect that future iterations of UHF fMRI could link layer-specific spinal cord responses to various aspects of human behaviour – addressing longstanding questions both within the domain of pain and beyond (e.g. touch and motor control).

Considering the spatial specificity achieved here, we envision that such mesoscopic fMRI approaches will act as a bridge (Finn et al., 2023), connecting human spinal cord research to that in other species. There, entire dorsal horn populations are now recorded (Greenspon et al., 2019; Shekhtmeyster, Carey, et al., 2023), but deeper layers have traditionally remained underexplored (Wercberger & Basbaum, 2019). Further advances are needed to establish the full potential of mesoscopic spinal UHF fMRI, but recent progress in coil design (Baskaran et al., 2025; Lopez-Rios et al., 2023), parallel transmit technology (Aigner et al., 2024; Papp et al., 2024), and correction for field inhomogeneities (Breheret et al., 2025; Vannesjo et al., 2019) will help here. This might furthermore extend our approach to the thoracic and lumbar spinal cord, promising to yield new insights into spinal control of internal organ and lower limb function. Ultimately, combining mesoscopic spinal imaging with concurrent behavioural and brain recordings (Tinnermann et al., 2021) as well as paradigms targeting distinct aspects of the complexities of bottom-up (Ding et al., 2025; L.-H. Wang et al., 2022) and top-down processing (Fatt et al., 2024; Heinricher et al., 2009), will allow for rich insights into the fundamental building blocks of nociception and the experience of pain.

## 4. Methods

### 4.1. Participants

We acquired data from four cohorts of healthy volunteers across *screening*, *discovery*, *validation* and *behavioural* datasets (Figure 1D). The *screening* dataset (Dataset 1: N = 62, 30 female, mean age: 27.6 years, range: 18-41 years) was used to select participants with adequate data quality based on anatomical and functional MRI scans without the use of any paradigm. The *discovery* dataset (Dataset 2; N = 16, 6 female, mean age: 31.3 years, range: 21 – 38 years) consisted of participants who were selected from the screening dataset based on the quality of their MRI data and from whom we acquired fMRI data using a robust pain paradigm. The preregistered *validation* dataset (Dataset 3; N = 25, 8 female, mean age: 30.4 years, range: 20 – 44 years) consisted of data acquisitions with an optimized fMRI protocol that made participant pre-selection unnecessary; note that the independent Dataset 3 employed the same pain paradigm as used in Dataset 2. The sample size for Dataset 3 was based on a preregistration (https://osf.io/pb6de) in which we defined a minimum target number of 17 participants to reach a pooled dataset size of 33 participants (Datasets 2 and 3 combined). This calculation was based on the upper bound of a bootstrapping approach contained in a recent 7T spinal cord fMRI study (Seifert et al., 2024), where it was estimated that a sample size of 23 to 33 participants would be necessary to detect a significant effect. Due to the availability of additional resources, we were able to exceed this minimum sample size and thus decided to include more participants, as this should allow to also detect significant effects in the *validation* dataset on its own. Please note that i) we aimed for sex-balanced samples but were restricted by the participant pool cleared for 7T MRI and volunteers’ willingness to participate in pain experiments and ii) due to a technical error, two participants were part of both Dataset 2 and 3, leaving 39 unique participants in the pooled dataset. Additionally, we acquired a *behavioural* dataset (Dataset 4: N = 20, 12 female, mean age: 30.1 years, range: 20-40), where we employed the same pain paradigm and obtained continuous pain ratings as well as various autonomic nervous system measures. Across all datasets, participants gave written informed consent. All studies were approved by the Ethics Committee of the Medical Faculty of the University of Leipzig.

### 4.2. MRI data acquisition

In the following, we list commonalities across data acquisition for Datasets 1-3. MRI data were acquired on a 7 Tesla MAGNETOM Terra scanner (Siemens Healthineers, Forchheim, Germany), equipped with a custom-built 24-receive-channel neck coil for cervical spinal cord imaging (Figure 1A-B; MRI.TOOLS, Berlin, Germany). The coil was designed for imaging of the entire cervical spinal cord (+ lower brainstem) with the contours conforming to the curvature of the neck. 16 receive channels are arranged in three rows along the back and 4 channels are positioned on either side of the neck and head. On the RF transmit side, each channel’s phase is optimized regarding efficiency (B1^+^ / specific absorption rate [SAR]) and groups of 4 channels are combined via power splitters, leading to 6 transmit channels (theoretically allowing for pTx functionality), which we used in single transmit mode by combining them via an additional 6-way power splitter.

To increase B_0_ homogeneity in the neck region (as well as to reduce bulk movement), we used a passive shimming approach by placing pads filled with a susceptibility-matched material (liquid perfluorocarbon; Sat Pad Inc., West Chester, USA) around the neck and onto the chest. At lower field strengths, similar implementations have been demonstrated for anatomic neck imaging (Cox & Dillon, 1995; G. Lee et al., 2015) and a number of 3T spinal fMRI studies have employed this technique (Weber et al., 2016a, 2016b), yet without evaluating its effects.

To mitigate the detrimental influence of physiological noise, which is prominent in spinal cord fMRI (Brooks et al., 2008; Cohen-Adad et al., 2010; Eippert et al., 2017; Piché et al., 2009; Verma & Cohen-Adad, 2014) and even more so at ultra-high fields (Vannesjo et al., 2018), we recorded cardiac and respiratory data in Datasets 2 and 3, allowing for extensive retrospective physiological noise correction of 7T fMRI data. Specifically, we measured respiratory activity via a breathing belt (Brain Products, Gilching, Germany) – and in Dataset 3 additionally via a breathing patch (BIOPAC Systems Inc., Goleta, CA, USA) – and cardiac activity via a pulse oximeter (Dataset 2: vendor-based product; Dataset 3: BIOPAC Systems Inc.).

To minimize cervical lordosis for all acquisitions, participants were instructed to tilt their head slightly towards their chest (Cohen-Adad et al., 2021). Participants were also instructed to 1) minimize head and body motion, 2) stay vigilant, 3) avoid excessive swallowing, and 4) avoid excessively deep breathing. The isocenter was set approximately to the participants’ larynx.

#### 4.2.1. Screening dataset (Dataset 1)

The anatomical and functional scans acquired in this dataset were part of a larger evaluation of different scan parameters for protocol optimization. Here, we only describe the anatomical scans and the functional scan we used to assess data quality, as necessary to select participants for Dataset 2.

##### Anatomical data

A high resolution T1-weighted image was acquired using a 3D gradient-echo VIBE sequence, employing the following parameters: 176 sagittal slices; resolution: 0.8 × 0.8 × 0.8 mm^3^; field of view: 220 × 220 mm^2^; TE: 1.56 ms; flip angle: 25°; TR: 10 ms. High resolution T2*-weighted images were acquired with 5 different echo times (4.55 ms, 8.12 ms, 11.65 ms, 15.17 ms, 18.69 ms), employing the following parameters: 15 transversal slices; resolution: 0.38 × 0.38 × 3.00 mm^3^; field of view: 146 × 137 mm^2^; flip angle: 45°; TR: 0.654 s; we acquired 3 runs of 5 echoes each.

##### Functional data

For each participant we acquired 100 volumes of BOLD data using a gradient-echo EPI sequence (https://www.cmrr.umn.edu/multiband/; Moeller et al., 2010) covering spinal cord segments C5-C7. EPI volumes were acquired with the following parameters: slice orientation: transverse oblique; number of slices: 16; slice thickness: 3.0 mm without gap; in-plane resolution: 0.75 × 0.75 mm^2^; TR: 1106 ms; TE: 23 ms; flip angle: 61°; field of view: 128 × 126 mm^2^; field of view position: centred on spinal cord between 3rd and 4th spinal discs (i.e., middle of C5 vertebra); GRAPPA acceleration factor: 3; partial Fourier factor: 6/8; phase-encoding direction: anterior-to-posterior (AP); echo spacing: 1.01 ms; bandwidth: 1130 Hz/Pixel; slice angulation: individually tilted, such that slices were perpendicular to the spinal cord in the area of interest. B_0_ shimming was performed using an adjust volume that was centred on the spinal cord in left – right and anterior – posterior directions, had a volume of 40 x 60 x 60 mm and was oriented the same way as the EPI slice stack.

#### 4.2.2. Discovery dataset (Dataset 2)

##### Anatomical data

A high resolution T1-weighted image was acquired with the same parameters as in the screening dataset. A high resolution T2*-weighted image was acquired with the same parameters as in the screening dataset, except for a reduced slice number (15). It was furthermore recorded with an additional echo, called navigator, that was acquired reading out a single line through the centre of k-space after the last echo in each TR (“phase stabilization” option in the vendor-provided GRE sequence).

#### Functional data

For each participant we acquired four runs of BOLD data (with 460 volumes each) using the same parameters as for the screening dataset, except for a reduced slice number (15) and a changed TR (1120 ms).

#### 4.2.3. Validation dataset (Dataset 3)

##### Transmit adjustment

Due to RF inhomogeneities along the cord, the transmit coil reference voltage was adjusted manually for each participant to achieve optimal flip angles inside the spinal cord at the segment of interest. To this end, a B1^+^ map was acquired with a pre-saturated 2D turbo-FLASH sequence with the following parameters: 56 sagittal slices (2 slice groups), resolution: 2.5 mm isotropic, field of view: 200 × 120 mm^2^, TE: 2.27 ms, TR: 34 s, flip angle: 10°, total acquisition time: 1:10 min. The ratio of the signal intensities of the images acquired with and without pre-saturation (90°) gives the cosine of the true flip angle voxel-wise (B1^+^ map). A region of interest (ROI) was drawn inside the spinal cord at the level of segment C6, and the mean flip angle measured within the ROI was used to calculate the optimal reference voltage required to obtain the target flip angle (90°). In cases where the optimal reference voltage exceeded the maximum allowable for the coil, the reference voltage was set to the maximum value and the flip angle was adapted accordingly.

##### Anatomical data

High resolution T1-weighted images were acquired using a 3D coronal MP2RAGE sequence (Marques et al., 2010) with 2 inversion times, from which the second inversion image was used. The sequence used in this work is a modified version of the product sequence, whose inversion pulse (hyperbolic secant) was changed (HS1 instead of HS4) to minimize SAR limitations potentially encountered in the spinal cord. Parameters were set as previously reported (Massire et al., 2016): resolution: 0.7 × 0.7 × 0.7 mm^3^; TE: 2.15 ms; flip angle: 4 / 5°; inversion time: 700/2400 ms; TR: 5 s.

A high resolution 2D T2*-weighted image was acquired using the same protocol as used in Datasets 1 and 2 (again including phase stabilization). To allow for an optimized offline reconstruction of the T2* acquisitions (minimizing the breathing artifacts), i) MRI data were linked to the respiratory recordings by including volume triggers and ii) a fully sampled, low-resolution GRE reference scan was acquired using the same slice geometry, with the following parameters: in-plane resolution: 2×2 mm^2^, TE: 3.16 ms, TR: 400 ms, single average.

##### Functional data

For each participant we acquired four runs of BOLD data (with 460 volumes each) using a single-shot gradient-echo EPI sequence covering spinal cord segments C5-C7. EPI volumes were acquired with the following parameters: slice orientation: transverse oblique; number of slices: 16; slice thickness: 3 mm without gap; in-plane resolution: 0.8 × 0.8 mm^2^; TR: 1123 ms; TE: 27 ms; field of view: 128 × 128 mm^2^; field of view position: centred on spinal cord between 3rd and 4th spinal discs; GRAPPA acceleration factor: 3; partial Fourier factor: 6/8; echo spacing: 1.02 ms; bandwidth: 1115 Hz/Pixel; slice angulation: individually tilted, such that slices were perpendicular to the spinal cord in the area of interest.

All EPI data were acquired using a custom-built sequence with a novel implementation of slice-specific z-shimming (optimized for 7T) to reduce signal loss and across slice variability (Eippert et al., 2024; Finsterbusch et al., 2012; Kaptan et al., 2022). Compared to earlier investigations at 3T, the slices in this study are thinner, reducing the EPI data’s vulnerability to signal loss. In contrast, susceptibility-induced B_0_ inhomogeneities are more pronounced at 7T, suggesting that z-shimming could offer significant improvements in signal recovery. The selection of slice-specific z-shims was carried out in an automated manner, using a modification of a previously developed procedure (Kaptan et al., 2022) based on the acquisition of a z-shim reference scan with 31 equidistant z-shim settings compensating gradients of 0.3 mT/m. Within-scan GRAPPA reference data for the EPI reconstruction were obtained with a FLASH-based acquisition which provided a significantly better performance compared to the standard EPI reference data in pilot measurements, in line with previous experiments (Seifert & Xu, 2022).

To minimize distortions in EPI data on a participant-by-participant basis, we chose a phase-encoding direction of either A-P or P-A, based on acquiring 10 EPI volumes with opposing direction and visually comparing the level of distortion in phase-encoding direction.

### 4.3. Thermal stimulation

In Datasets 2, 3 and 4 we employed painful heat stimuli (30 s duration), which were applied via an MR-compatible thermode with a ramp-speed of 70°C/s and a circular stimulation zone with a diameter of 27 mm (PATHWAY CHEPS; Medoc, Ramat Yishai, Israel). The stimuli were administered to the inner left forearm of the participants, most likely corresponding to dermatome C6 (M. W. L. Lee et al., 2008). In order to minimize possible sensitization and habituation over runs, we changed the stimulation area (usually after two of the four runs), ensuring that different skin regions within the middle of the forearm were used. Changing the thermode position in the MRI environment was realized by a custom-built MR-compatible extension mechanism (Müller et al., 2024). This was attached to the thermode to allow for easy repositioning between runs from outside the scanner bore, without having to move the scanner table on which participants were lying. Each stimulus consisted of an individually determined plateau temperature between 41°C and 45°C and thirty pseudo-randomized heat spikes of 2, 3, or 4 degrees superimposed on the plateau, each lasting less than a second (Figure 1C). This stimulation protocol was based on previous studies (Brooks et al., 2017; Oliva et al., 2021, 2022) and was chosen to maintain a painful percept throughout stimulation, while at the same time avoiding sensitization and skin damage (Lautenbacher et al., 1995). The baseline thermode temperature was set to 32°C.

### 4.4. Experimental procedure

At the beginning of the experiment, participants were informed about the study and any remaining questions were discussed. For Datasets 2, 3 and 4, we additionally explained the procedure of the pain experiment and before the main experiment started, participants were familiarized with the heat stimuli. Starting with a plateau temperature of 40°C (with spikes up to 44°C), the stimuli were increased adaptively in 0.5 or 1°C steps and participants were asked to rate each 30 s stimulus on a numerical rating scale (NRS) with the following anchors: 0 = no sensation, 50 = pain threshold, and 100 = unbearable pain. We aimed for a target temperature corresponding to a rating of ∼75 on the NRS (i.e., clearly in the painful range while being bearable), but set a plateau maximum of 45°C (with spikes up to 49°C) as this was the highest temperature allowed for safety reasons. The average plateau temperature used was 43.2°C (range: 41-45°C) in Dataset 2, 43.4°C (range 42-45°C) in Dataset 3 and 42.8°C (range 41-45°C) in Dataset 4.

During the experiment, participants were informed about the start of the functional runs. The thermal stimuli were then applied in an On-Off block design in which 30 s rest periods were followed by 30 s heat pain periods. Overall, the experiment involved four runs with eight pain stimuli each (Figure 1C).

During the behavioural experiment (Dataset 4), participants were additionally asked to continuously rate their pain experience. In alternating runs, participants rated either the intensity or the unpleasantness of the pain stimuli, with the starting condition randomized. The distinction was explained to the participants using a radio metaphor, similar to previous investigations (Nahman-Averbuch et al., 2023; Price et al., 1983; Rainville et al., 1992). The scale had the anchors “not at all intense/unpleasant” and “as intense/unpleasant as imaginable”. In the MRI environment (Datasets 2 and 3) participants were only asked after each functional run to verbally rate the stimuli again (using the NRS previously used for calibration), as providing ratings during the scan would have resulted in confounding sensorimotor activation in the cervical spinal cord and was thus not feasible. In case stimuli were experienced as too painful or not painful anymore, the temperature was adapted in steps of 0.5 or 1°C for the subsequent run. Necessary adaptations included both lowering of temperatures (Dataset 2: 2, Dataset 3: 2, Dataset 4: 8 times) and increasing of temperatures (Dataset 2: 7, Dataset 3: 10; Dataset 4: 4 times). Based on this rating, we excluded two runs from one participant in Dataset 2 and two runs in three participants each in Dataset 3, as they were clearly not painful for the participants (see section 4.6). The remaining runs contributed to an average rating across all runs that was 68.8 (range 52-88) in Dataset 2 and 66.2 (range 55-77.5) in Dataset 3, thus slightly below our target of NRS-75, but clearly in the painful range (i.e. above NRS-50).

### 4.5. Additional recordings

In Dataset 2, we additionally attached two Ag/AgCl electrodes filled with isotonic electrolyte gel to the thenar and hypothenar eminences of the left hand to record skin conductance responses (SCR). The cardiac data recorded for physiological noise correction in both Dataset 2 and 3 was also used to assess stimulus-evoked heart period responses (HPR).

In Dataset 4 we again employed these previously used methods to assess stimulus-locked SCR and HPR using a BrainAmp ExG system (Brain Products, Gilching, Germany). Cardiac recordings were realized here via bipolar electrodes placed on the right sternum and below the rib cage to the left side of the participant’s body, with the ground electrode placed close to the rib electrode. Since this dataset was acquired outside the MRI environment, we were able to gather additional pupil diameter data using an EyeLink 1000 system (SR Research, Ottawa, ON, Canada) with a sampling rate of 1 kHz.

### 4.6. Data exclusion

In Dataset 1, we did not record functional data for three participants due to severe artifacts (severe shim problems in two participants, high frequency spikes in another participant), leaving N = 59 for the analyses. In Dataset 2, we excluded data from two runs in one participant, as the thermode had not been optimally attached during these runs and the participant reported the stimulation not to be painful or not even noticeable. In Dataset 3 we excluded data for the same reason from two runs in three participants each. Additionally, in Dataset 3 for one participant, we were only able to record three runs as their hand became numb. As stated in the preregistration (https://osf.io/pb6de), we included all participants whose data encompassed at least 2 out of 4 runs (thus retaining 97% of functional data in Dataset 2 and 93% in Dataset 3). For one participant, we were not able to acquire the T2*-weighted image, as they asked to end the experiment after the functional runs due to not being comfortable in the scanner anymore. This participant is excluded in the analysis focusing on individual data where the T2*-weighted data are necessary (Figure 4D-F) but is included in all other analyses. As mentioned above, due to a technical error two participants were part of both Datasets 2 and 3, leaving 39 unique participants in the pooled dataset. To avoid data duplication, we used their data from Dataset 3 and excluded their data from Dataset 2 in these pooled dataset analyses.

### 4.7. FMRI data analysis

Preprocessing and statistical analyses were carried out using FMRIB Software Library (FSL, version 6.0.3, https://fsl.fmrib.ox.ac.uk/fsl/, Jenkinson et al., 2012), Spinal Cord Toolbox (SCT, version 6.1, https://spinalcordtoolbox.com, De Leener et al., 2017), Advanced Normalization Tools (ANTs, version 0.0.8, http://stnava.github.io/ANTs/, Avants et al., 2009) as well as custom Bash and Python (version 3.9) scripts.

#### 4.7.1. Preprocessing

##### 4.7.1.1. Thermal denoising

As a first processing step for all datasets, we applied Marchenko-Pastur principal component analysis (MP-PCA, https://github.com/NYU-DiffusionMRI/mppca_denoise, Veraart et al., 2016) to the raw EPI data to reduce thermal noise (Ades-Aron et al., 2021; Kaptan et al., 2023). The procedure makes use of the assumption that noise-related eigenvalues follow a Marchenko-Pastur distribution and can thus be distinguished from signal-related eigenvalues.

##### 4.7.1.2. Motion correction

For all datasets, motion correction was carried out in two steps. First, we created a mean image of the first run’s EPI volumes after thermal noise correction via MP-PCA. This image was segmented in order to create a cord-centred cylindrical mask, which was used to prevent adverse effects of non-spinal movement on the motion parameter estimation. Motion correction to this target image was then carried out slice-wise (allowing for x- and y-translations), using spline interpolation and a 2^nd^ degree polynomial function for regularization along the z-direction (De Leener et al., 2017). Second, we repeated the motion correction of the original time series, now employing the mean image of the initially motion-corrected time-series of the first run as the target image. The reason for this two-stepped procedure is that at 7T, apparent motion artifacts are more severe than at lower field strengths, and the first uncorrected mean image is rather blurry and thus a suboptimal choice for a target image. To demonstrate this, we assessed the sharpness of the two target images by determining the Laplacian variance and observed that the target image in the area of the motion-correction mask is improved by 10.6% in Dataset 1, 18.9% in Dataset 2 and 16.5% in Dataset 3 when using a second step.

##### 4.7.1.3. Participant selection process for discovery dataset

Selection of participants from the screening dataset (Dataset 1) for inclusion in the discovery dataset (Dataset 2) was based on three factors and occurred after data were corrected for thermal noise and motion (sections 4.7.1.1 and 4.7.1.2). First, we empirically assessed participants’ EPI data quality by investigating the similarity of functional and anatomical images in terms of the spinal cord’s cross-sectional shape. This was necessary, as at 7T, spinal cord EPI data can suffer strongly from signal loss and distortion (Figure 2C), which are not similarly present in the ME-GRE data (thus presenting a ground-truth). Since the severity of these artifacts is due to individual differences in anatomy, they are reproducible over sessions and thus allow to select participants with adequate data quality for Dataset 2. To assess similarity for each participant, we created a cord mask in both the functional images and the ME-GRE images (see 4.7.1.8) and calculated the Dice coefficient of these two masks (Figure 2D), with higher values indicating stronger similarity. Second, we conducted a visual assessment of the EPI images, with a particular focus on ensuring that data in target segment C6 was of high quality. Third, recruitment of participants for Dataset 2 also depended on their willingness to take part in a pain experiment as well as their availability during the data acquisition period; for these reasons, the participant with the highest Dice coefficient is for example not included in Dataset 2.

##### 4.7.1.4. Motion in z-direction

Due to the increased SNR and resolution in z-direction (i.e., slice thickness of 3 mm), we were able to detect slight movements in the z-direction in Datasets 2 and 3 (which, in this project, were likely introduced by changing the thermode position between runs). To address this, we developed a general-purpose algorithm to perform translations in the z-direction before finally applying the previously described motion correction procedure (section 4.7.1.2), ensuring that these subtle movements were accurately captured, because they would have been missed by the slice-wise in-plane motion correction algorithm.

To estimate the necessary z-translation, we determined how much the discs’ rostro-caudal location of each run changed in comparison to the first run’s mean image. We first moved a cord segmentation mask by an individually determined amount onto the part of the image where the discs are located. Both the mask and the motion-corrected mean EPI image were up-sampled by a factor of four using Non-Local MRI Upsampling (Manjón, Coupé, Buades, et al., 2010), i.e. a super-resolution approach that uses a data-adaptive patch-based reconstruction. A disc image was created by restricting the mean EPI image to the area of the discs. To enhance the contrast and to optimize the algorithm, we restricted the image to a specific value range and dilated the disc images by 7 voxels within each slice. As a main step, we then aligned the disc image of each run with the first run using antsRegistration, restricting the deformation to z-direction. The resulting translation is described by one number which is then down-sampled by the chosen factor of four. This down-sampled transformation is used to shift the original denoised images to the correct position in z-direction using antsApplyTransform. These z-motion corrected data are then motion-corrected for the remaining in-plane motion, now aligning the correct slices.

We only used this step when motion in z-direction in a run was greater than or equal to 0.75 mm compared to the first run (i.e., ¼ of the slice thickness). This was the case for one run in one participant and two runs in another participant in Dataset 2 (three corrections overall) and one run in two participants and two runs in three participants in Dataset 3 (eight corrections overall).

##### 4.7.1.5. Physiological noise correction

In Datasets 2 and 3 we used the concurrently recorded physiological data to address physiological noise in EPI data during general linear model (GLM) estimation. In the cardiac data, R-peaks were automatically detected (as well as manually checked) and intervals with poor cardiac data quality were marked to be scrubbed during model estimation. The respiratory data were linearly detrended and z-scored. Please note that for Dataset 3 (due to equipment instability), we recorded respiration both with a pad and a belt and mostly used the pad data except when these data exhibited artefacts.

Based on the cardiac peaks and normalized respiration data, we employed a physiological noise model (PNM) to address fMRI signal variations attributed to cardiac and respiratory activities. This model, adapted from the RETROICOR method (Glover et al., 2000), calculates the cardiac and respiratory phases at which the slices were captured. Specifically tailored for spinal cord fMRI, this approach generates slice-specific physiological regressors for further analysis (Brooks et al., 2008; Kong et al., 2012). By employing Fourier basis series with sine and cosine harmonics, we derived 8 noise regressors each for cardiac and respiratory influences, and 16 more for their interaction effects, culminating in a total of 32 noise regressors.

Additionally, we used PNM to model the cerebrospinal fluid (CSF) signal as another regressor by extracting the signal from voxels within a CSF mask that exhibit high levels of signal variance. For the CSF mask creation, we used the output of the function sct_propseg, dilated this mask by 3 voxels and subtracted a manually corrected cord mask. We refrained from using respiratory volume per time (RVT) as additional regressor because this led to rank-deficient design matrices when used in combination with the extraspinal PCA regressors (described in next section).

##### 4.7.1.6. Additional noise correction

In addition to PNM, we also added further noise regressors to the first-level GLM to remove residual motion and physiological noise. In spinal cord EPI data, (apparent) motion artifacts arise from several sources, such as bulk movement, respiration or swallowing (Kinany et al., 2023). First, we used the estimated slice-specific motion parameters in x and y direction to create higher-order motion parameters (i.e., original x- and y parameters, temporal derivative of x- and y-parameters (calculated backwards), squared x- and y-parameters and squared temporal derivative of x- and y-parameters). Especially datasets with greater degrees of motion are likely to benefit from this high-parameter approach (Satterthwaite et al., 2013). Second, we identified volumes with excessive motion remaining after motion-correction by calculating the root mean square intensity difference between successive volumes (dVARS) as well as the root mean square intensity difference of each volume to a reference volume (refRMS) using FSL’s fsl_motion_outlier algorithm. Volumes presenting with dVARS or refRMS values two standard deviations above the time series mean were defined as outliers and individually modelled as regressors of no interest in the first level GLM (on average, 3.3% of volumes (15.3 ± 8.2 volumes of the 460 volumes per run) were classified as outliers). Third, we defined a region around the spinal cord as an “extra-spinal noise region” (created by dilating the cord mask by 18 voxels and then subtracting the cord mask from the dilated mask, creating a ring-like structure). Following earlier work (Hemmerling et al., 2023, 2025) we performed a slice-wise principal component analysis (PCA) in that region and included the first 5 components from this PCA as additional slice-wise noise regressors in the first-level GLM.

##### 4.7.1.7. Cord segmentation

In order to obtain a high-resolution segmentation of the spinal cord, we used the T1-weighted acquisitions acquired in each dataset (resolution: 0.8 mm (Dataset 1 and 2) or 0.7 mm (Dataset 3) isotropic voxels). We first cropped the images in the z direction from above vertebra C1 to below T1 based on an automated vertebral labeling step. Since the MP2RAGE images suffered from aliasing artifacts, we also cropped all images from both datasets in x and y direction to a box of size 100 × 120 voxels. We then applied ANT’s N4 bias field correction algorithm on the cropped structural image to correct for intensity inhomogeneities (Tustison & Gee, 2010). As a next step, we denoised the structural data via Adaptive Optimized Nonlocal Means filtering (Manjón, Coupé, Martí-Bonmatí, et al., 2010) to account for spatially varying noise levels and to increase the SNR. We then segmented the bias-corrected and denoised T1-weighted data in an two-step procedure to improve the robustness and quality of the final segmentation: the data were initially segmented using the SCT DeepSeg algorithm (Gros et al., 2019), were then smoothed along the spinal cord centerline using an anisotropic Gaussian kernel with 6 mm sigma, and again segmented via the DeepSeg algorithm.

To obtain a segmentation of the spinal cord in the EPI data, we used the mean image of the MP-PCA enhanced and motion-corrected time series as input for SCT’s DeepSeg algorithm. All EPI-based segmentations were visually inspected and manually corrected if necessary.

##### 4.7.1.8. Grey matter segmentation

To obtain robust and precise grey matter segmentations for participant-specific analyses of dorsal horn BOLD responses in Datasets 2 and 3, we used the T2*-weighted ME-GRE images, which were acquired for this purpose with very high in-plane resolution (0.38 mm) and an additional navigator. After the acquisition of Dataset 2, a new method for navigator-based correction became available (Beghini et al., 2025), which we applied to Dataset 3 (note that this could not be applied retrospectively to Dataset 2, as it required the k-space data to be saved during data acquisition).

This method (developed specifically for the spinal cord) was applied to the raw data using MRINavigator.jl (v0.0.1, https://github.com/NordicMRspine/MRINavigator.jl, Julia v 1.8.5) and uses the navigator phase to measure and correct for breathing-induced B_0_ field fluctuations (Beghini et al., 2025). The field fluctuations cause ghosting and signal loss in the images and are especially intense in the spinal cord due to the proximity of the lungs. The FFT navigator processing approach from MRINavigator.jl was used, in which the navigator profiles first undergo a 1D Fourier transform after which a region of interest (4cm centred on the spinal cord) is selected and used to estimate the phase. When phase wrapping occurred, an additional unwrapping step was included in the processing pipeline (‘FFT unwrap’ from MRINavigator.jl). The unwrapping algorithm makes use of the respiratory belt recordings, which were synchronized with the raw MR data using the PhysIO toolbox (Kasper et al., 2017). In rare cases, the unwrapping step produced irregular phase estimates, and the processing was reverted to the original FFT navigator approach. The resulting phase estimates were used to remove the corresponding phase from the imaging data by demodulation. After the navigator-based correction, the images were reconstructed offline with a conjugate-gradient SENSE algorithm (10 iterations, L2 regularization; Knopp & Grosser, 2021; Pruessmann et al., 1999). The sensitivity maps for the reconstruction were calculated from the GRE reference scan using the ESPiRIT algorithm (Knopp & Grosser, 2021; Uecker et al., 2014), and were masked using MRINavigator.jl. A noise pre-whitening step, aimed at reducing the noise correlation across coils was applied before the reconstruction for SNR optimization purposes (Pruessmann et al., 2001).

For all three datasets, ME-GRE data were acquired with 5 echoes and three repetitions. The root-mean-square (RMS) image over the five echoes was computed for each run and then motion correction of the three runs was performed to obtain one motion-corrected mean image for each participant.

ME-GRE images served two purposes. First, for participant selection for Dataset 2 (see section 4.7.1.3), ME-GRE data were needed to assess the similarity between anatomical (ME-GRE) and functional (EPI) data. For this, we used SCT’s deepseg function to create a cord mask from the ME-GRE data, that was then registered to the mask of the motion-corrected functional mean image (via slice-wise translations in x- and y-direction), after which similarity could be assessed. Second, ME-GRE data were needed for precise grey matter segmentation as employed in the analysis of participant-level BOLD responses (Datasets 2 & 3). For this endeavour, we used SCT’s deepseg_gm function to create a grey matter mask, that was registered to the motion-corrected mean image space in a segmentation-based transformation step using a registration based on B-Spline regularized non-linear symmetric normalization. To delineate the left and right dorsal and ventral horns from this overall grey matter mask in each individual (Figure 4D), we performed the following steps in an automated fashion. First, we identified the bounding box surrounding the mask and determined the midpoint in the left/right direction. Second, we traced along the dorsoventral line through this midpoint, starting from the dorsal horns’ end, to locate the first point with data in the grey matter mask, identifying it as the central part of the grey matter (i.e., the grey commissure). Third, we defined this as the midpoint between the ventral and dorsal regions. Finally, we created four grey matter horn masks based on these lines, deliberately leaving the boundary line between the left and right masks unmarked.

##### 4.7.1.9. Registration to template space

Individual-level statistical analyses were conducted within each participant’s native space, while group-level analyses were performed in a standardized space defined by the PAM50 spinal cord template (De Leener et al., 2018). To transform individual data from native to template space, we used the T1-weighted image that had been corrected for bias and noise. Vertebral levels were identified and labeled, and the spinal cord was straightened. The structural image was then registered to the template using an iterative, non-rigid registration method (De Leener et al., 2017). The initial step of this method relied on segmentations, while the second step involved either a segmentation or an image-based registration, with the optimal approach chosen for each participant. The resulting inverse warping field was used to initiate the registration of the PAM50 template to the motion-corrected mean functional image through SCT’s multi-step non-rigid registration based on segmentations (De Leener et al., 2017). This registration provided a warping field that allowed us to transform relevant images from each individual’s native space to template space using spline interpolation. We applied this warping field to i) the mean functional image, ii) the tSNR image and iii) the statistical output from GLM estimation (i.e. parameter estimate images).

#### 4.7.2. Statistical analysis

##### 4.7.2.1. General linear model

Statistical analysis of the fMRI data was carried out using tools from FSL. At the level of individual runs, a general linear model (GLM) approach implemented in FSL’s FEAT (FMRI Expert Analysis Tool; http://fsl.fmrib.ox.ac.uk/fsl/fslwiki/FEAT; Woolrich et al., 2001) was employed, including spatial smoothing with an isotropic 2 mm FWHM Gaussian kernel (via SUSAN), high-pass filtering at 100 s and correction for temporal autocorrelation (via FILM). Three different GLMs were created.

The first (and main) GLM’s first-level design matrix consisted of a regressor over the whole duration of the 30 s heat stimulus convolved with a double-gamma hemodynamic response function (HRF), thus assessing a typical block-response. The second GLM aimed at assessing the reliability of the 30 s heat response. For that purpose, we performed an odd-even split by creating a GLM that contained one regressor for all odd trials and one regressor for all even trials. The third GLM focused on an onset response to the heat stimulus by modelling the first 3 s of the heat stimulus as our task-based regressor of interest (the remaining 27 s were also modelled but not investigated). This analysis was theoretically motivated, as it has been demonstrated that the input signals received by the spinal cord (action potentials in peripheral nerves) exhibit two different types of responses to 30 s heat stimuli: A-δ fibres in primates show a ‘sustained’ type I response that gradually increases over time and a ‘phasic’ type II response with an early peak that quickly diminishes after about 3s (Treede et al., 1995, 1998); slow and quick responses have also been distinguished in C fibres (Johanek et al., 2008; Meyer & Campbell, 1981). Since this was an exploratory analysis in Dataset 2, we preregistered this analysis before acquiring Dataset 3 to test whether such a pattern would emerge in this replication dataset (https://osf.io/pb6de).

The following regressors of no interest were added to all of the above design matrices for robust denoising: 33 slice-wise PNM regressors, eight slice-wise motion regressors, five slice-wise extra-spinal PCA regressors and one regressor for each volume with excessive motion.

##### 4.7.2.2. Vascular autorescaling (VasA)

The statistical power of fMRI group studies is inherently limited if participants differ in their baseline physiology (e.g., blood volume or baseline deoxyhemoglobin concentration). To account for such vascularization differences between participants, we used a vascular autocalibration method that has been physiologically validated and shown to increase the functional sensitivity in group analyses of brain fMRI data at 7T (Kazan et al., 2016, 2017) without introducing bias. VasA estimates the local vascular reactivity from the residuals of the fMRI time-series after regressing out the task components, and renormalizes the local fMRI amplitude estimates based on the VasA estimate. We therefore used this method for all group analyses; note that VasA only adjusts for inter-individual differences and thus has no effect on single-participant analyses.

##### 4.7.2.3. Group analysis heat response

After obtaining VasA-corrected β-estimate maps (i.e., COPE images in FSL) for each run, we combined these in a fixed-effect participant-level analysis using FSL’s FEAT. The resulting participant-wise COPE images were brought to template space using the previously obtained warping fields. For all three GLMs, we submitted the template space COPE images to a one-sample t-test to examine whether the heat stimulus evoked a significant BOLD response in our hypothesized target region (left dorsal horn C6). Stringent correction for multiple comparisons was carried out via non-parametric permutation testing in our target region as implemented in FSL’s randomise algorithm (Winkler et al., 2014) using Threshold-Free Cluster Enhancement (TFCE; Smith & Nichols, 2009) and controlling the family-wise error rate (FWE). We also present uncorrected results for the whole cord area to describe the spatial distribution of the activation in an unbiased manner. We present all analyses for Dataset 2 and 3 separately, as well as for the combined dataset.

### 4.8. Atlas creation

In order to capitalize on the improved resolution at ultra-high field strength and to thus be able to describe the activation patterns in more fine-grained detail, we created a grey matter atlas of the human spinal cord based on Rexed’s laminae (Rexed, 1952, 1954). This laminar organization of the spinal cord was originally identified in cats, but is present across mammals (Tan et al., 2023), including humans (Schoenen & Faull, 2004) and is currently the major cytoarchitectonic organization scheme of the spinal cord (but see also newer approaches, e.g. Häring et al., 2018).

This atlas is based on a cross-section of a post-mortem human spinal cord at segment C6, from which histological analyses allowed for the creation of a grey matter atlas with labelled laminar divisions (as shown in “Atlas of the Spinal Cord”; Sengul et al., 2013); note that this atlas is complementary to the recently presented AMU7T atlas (https://github.com/spinalcordtoolbox/template_AMU7T; Le Troter et al., 2023), which is however not based on a laminar organization. We created a digital version of this cross-section whereby we grouped laminae I and II into the ‘superficial layers of the dorsal horn’, laminae III and IV into the ‘middle layers of the dorsal horn’, laminae V and VI into the ‘deep layers of the dorsal horn’ and laminae VII-IX into the ‘ventral horn’ (midline lamina X was not included here). This grouping was based on the following reasoning: i) a distinction between individual layers (e.g. I vs II) is not achievable with the current resolution of the here-employed PAM50 space and was thus not attempted; ii) the grouping into superficial, middle and deep layers is based on typical characterizations of the dorsal horn; iii) the creation of a single ventral horn mask was chosen because the focus of this study is the dorsal horn and hence only this structure was subdivided. From this vector graphic, we created four high-resolution NIfTI-images (in-plane resolution: 0.1 mm × 0.1 mm; each corresponding to one layer-group) and registered them to the PAM50 space. For this registration, we followed several steps outlined in previous work where an atlas for white matter tracts was created (Lévy et al., 2015).

In detail, we first resampled the grey matter segmentation included in the PAM50 template to an in-plane resolution of 0.1 mm × 0.1 mm and binarized it at a threshold of 50%. A slice corresponding to the middle of segment C6 was extracted and used as a reference for registering the binary grey-matter mask of the atlas (thus incorporating all layers). The registration was performed in ANTs and used an initial affine transformation and a non-rigid deformation using the SyN algorithm (for parameters see Lévy et al., 2015). Then, we used our reference grey matter segmentation slice to register this to all other C6 slices of the upsampled PAM50 grey matter segmentation using B-Spline regularized non-linear symmetric normalization (BSplineSyN). This warping field was then concatenated with the previous transformation to warp the atlas labels to each slice of spinal segment C6. The resulting slice-wise atlas images thus matched with the PAM50 gray matter and were merged into one file per laminar group and down-sampled to the PAM50 resolution of 0.5 mm × 0.5 mm.

### 4.9. Analysis of peripheral physiological responses and continuous ratings

Finally, we analysed the stimulus-evoked peripheral physiological response and continuous ratings to assess what type of response profiles a dynamic 30 s heat stimulus as used here elicits at different processing levels. These data consist of skin conductance (Datasets 2 and 4), heart period responses (Datasets 2-4), pupil dilation responses (Dataset 4) and continuous ratings of intensity and unpleasantness (Dataset 4).

4.9.1. *Skin conductance responses*

Skin conductance data recorded in Datasets 2 and 4 were initially filtered with a bidirectional first-order Butterworth bandpass filter using cut-off frequencies of 1 Hz and 0.0159 Hz (corresponding to a time constant of 10 s) and then down-sampled to 100 Hz. Data were cut into epochs ranging from -5 s to 35 s relative to the beginning of the heat stimulus of each trial and were baseline-corrected (at trial-time 0 ms). To distinguish responders and non-responders, we tested whether a participant had at least two runs that contained responses in at least 25% of the trials (a response being defined as an amplitude increase of 0.01 μS in a response window of 1 s to 30 s relative to the beginning of the heat stimulus in comparison to an average baseline amplitude in a baseline window -1 to 0 s before this window (Dawson et al., 2016)). Based on this criterion, three participants in Dataset 2 were defined as non-responders, resulting in N = 13 for the SCR analyses in this dataset.

#### 4.9.2. Heart period responses

Cardiac data from Datasets 2, 3 and 4 were initially processed by detecting R-peaks using MNE-Python (https://mne.tools, Gramfort, 2013). Afterwards, a manual correction of R-peaks was performed using in-house functions. To transform the R peaks into a continuous signal we assigned each inter-beat-interval to its following heartbeat. The time series was linearly interpolated with a sampling rate of 10 Hz and additionally filtered with a 2nd-order Butterworth filter from 0.01 to 2 Hz (Paulus et al., 2016). Data were then cut into epochs ranging from -5 s to 35 s relative to the beginning of each heat stimulus and baseline-corrected by subtracting the value of stimulation onset.

#### 4.9.3. Pupil dilation responses

Eye tracking data recorded in Dataset 4 were initially processed using MNE-Python to linearly interpolate missing data due to blinks as detected by the EyeLink Software in an interval from 50 ms before to 200 ms after the blink event. Subsequently, pupil data were processed with in-house Python scripts utilizing MNE functionality. The data were filtered with a first-order low-pass Butterworth filter with a cut-off frequency of 2 Hz, and remaining blink artifacts were manually detected. Data were cut into epochs ranging from -5 s to 35 s relative to the beginning of each heat stimulus and were baseline-corrected (at trial-time 0 ms). Epochs that contained more than 50% interpolated data points were excluded from the analysis (0.16% of all data). To account for differences in measured pupil dilation due to factors like brightness in the room, we z-standardized all values in the epochs (i.e. within a participant) before creation of grand average plots.

#### 4.9.4. Continuous ratings

Whenever participants interacted with the buttons on the rating device, the scale was updated and new data points were generated. Otherwise, the current state of the scale was recorded as a data point every 20 milliseconds. As a result, the sampling rate varied slightly, necessitating interpolation of the samples before further processing. Data were then cut into epochs ranging from -5 s to 35 s relative to the beginning of each heat stimulus and were baseline-corrected (at trial-time 0 ms). One trial of one participant was excluded from the analysis because the participant did not provide a rating and later indicated that they had forgotten to do so.

#### 4.9.5. Cluster-based permutation tests

For skin conductance responses, pupil dilation responses and both types of continuous ratings we conducted cluster-based permutation tests (Maris & Oostenveld, 2007) to assess whether the average stimulus-locked time courses are larger than zero. We searched within a time window of 0 s to 30 s relative to the beginning of heat stimulation for clusters of data points that have a significant value of the test statistic. This procedure is repeated 10000 times with labels being assigned randomly to the data. Using the distribution resulting from those permutations as a null distribution, significance of the found clusters can be established. Note that this approach could not be applied to cardiac data due to the tri-phasic pattern of the stimulus-evoked responses.

### 4.10. Open Science

The experimental plan – including sample size rationale as well as preprocessing and analysis steps – was preregistered before acquisition of Dataset 3 and the preregistration is openly available on the Open Science Framework (https://osf.io/pb6de). Any differences between preregistration and manuscript are listed in Table S1. The underlying data of all four datasets are currently only accessible to reviewers, but will be made openly available upon publication via OpenNeuro (Dataset 1-3) and OSF (Dataset 4). All code to reproduce the here-reported results is publicly available via GitHub (https://github.com/eippertlab/spinalcord-fmri-7T). The intended data sharing was mentioned in the Informed Consent Form signed by the participants and approved by the Ethics Committee at the Medical Faculty of the University of Leipzig.

## Acknowledgements

We would like to thank all volunteers who participated in these studies. Additionally, we want to thank everyone who assisted in data acquisition, especially Mandy Jochemko, Domenica Klank, Anke Kummer and Sylvie Neubert for their help with the MRI measurements; and Helene Minkus, Johanna Seemann and Clara Wilken for their help with the behavioural measurements. We would further like to thank Jöran Lepsien and Roland Müller for their technical help and Lisa-Marie Pohle for her input on visualization.

## Author contributions

Author contributions are listed alphabetically according to CrediT taxonomy (https://credit.niso.org).

Conceptualization: F.E., U.H.

Data curation: U.H.

Formal analysis: L.B., F.E., U.H.

Funding acquisition: F.E.

Investigation: F.E., N.G-W., U.H., Y.R., R.T., S.J.V.

Methodology: V.C., A.D., F.E., J.F., N.G-W., U.H., M.K., S.J.V.

Project administration: F.E., U.H.

Resources: J.F., H.E.M., K.J.P., R.T., N.W.

Software: L.B., A.D., F.E., U.H., M.K.

Supervision: F.E., N.W.

Visualization: U.H.

Writing - original draft: F.E., U.H.

Writing - review and editing: L.B., V.C., A.D., F.E., J.F., N.G-W., U.H., M.K., H.E.M., K.J.P., Y.R., A.T., R.T., S.J.V., N.W.

## Funding

F.E. received funding from the Max Planck Society and the European Research Council (under the European Union’s Horizon 2020 Research and Innovation Program; grant agreement No 758974). N.W. received funding from the Deutsche Forschungsgemeinschaft (DFG, German Research Foundation) – project no. 347592254 (WE 5046/4-2); the Federal Ministry of Education and Research (BMBF) under support code 01ED2210; the European Union’s Horizon 2020 research and innovation programme under the grant agreement No 681094.

## Declaration of competing interest

The Max Planck Institute for Human Cognitive and Brain Sciences and Wellcome Centre for Human Neuroimaging have institutional research agreements with Siemens Healthineers. N.W. holds a patent on acquisition of MRI data during spoiler gradients (US 10,401,453 B2).

