## Supplementary Material for "Ultra-high-field fMRI reveals layer-specific responses in the human spinal cord"

### 1. Passive shimming investigations

To assess the effect of passive shimming on field homogeneity, we acquired an additional pilot data set (N=7; 5 female; mean age 29.9 year; age range 23 - 42 years) not further described in the main manuscript. Using a within-participant approach, we acquired data with and without placing pads filled with a susceptibility-matched material (SatPads) around participants' neck and on their chest; the data acquisition order was counter-balanced regarding acquisitions with or without passive shimming.

MRI data were acquired using the same scanner and coil. In each participant and condition, we acquired one T1-weighted image of the entire cervical spinal cord (3D VIBE, voxel size: 0.8×0.8×0.8 $mm^3$) and two field maps of the entire cervical spinal cord (voxel size: 2×2×2 $mm^3$): one field map using only the vendor provided tune-up shim and one field map using the vendor's active shim procedures.

We processed the field map data both with and without SatPads by first unwrapping them using ROMEO (https://github.com/korbinian90/ROMEO, Dymerska et al., 2021) and then converting them from Hertz (Hz) to parts per million (ppm). For the anatomical images, we segmented the spinal cord using SCT's deepseg function. Manual corrections were applied to the segmentation if needed, and the images were cropped between the C1 and C7 vertebrae. This cord mask was then utilized to extract the ppm signal from each field map.

Group data (N=7 for active shim; N=6 for tune-up shim due to one participant's data not being available because of a scanner operator error) showed a consistent decrease in field inhomogeneity in acquisitions with passive shimming compared to acquisitions without passive shimming (37% with tune-up shim; decrease in 6/6 participants; 23% with active shim; decrease in 5/7 participants). See also Figure 2B for field map data from one exemplary participant.

### 2. Results for individual datasets

#### a. Sustained response

In addition to the group-level analyses reported for the combined dataset, we tested whether we would be able to detect spinal cord BOLD responses in response to the 30s heat stimulation within each dataset (i.e., the *discovery* dataset [Dataset 2] and the *validation* dataset [Dataset 3]), using identical analysis parameters as in the combined dataset. At an exploratory threshold ($p < 0.01$), in both datasets we observed the most prominent activation in target segment C6 (Figure S1A, S1D), found primarily on the ipsilateral (left) side and in both dorsal and ventral horn, but also extending contralaterally (Figure S1C, S1F). Generally, more widespread activation was present in the higher-powered Dataset 3, also leading to higher statistical scores overall. As mentioned already in the preregistration, Dataset 2 responses (which provide the first 7T demonstration of sustained dorsal horn BOLD responses) did not survive strict statistical thresholding using permutation testing in the target region of the left dorsal horn in segment C6 (Figure S1B; peak voxel: $t = 2.83$, $p_{FWE} = 0.055$), but inspired the acquisition of Dataset 3. In Dataset 3 however, two clusters were observed in the target region of the left dorsal horn in segment C6 (Figure S1E; cluster 1: 27 voxels, peak voxel: $t = 3.58$, $p_{FWE} = 0.006$; cluster 2: 3 voxels, peak voxel: $t = 3.50$, $p_{FWE} = 0.026$).

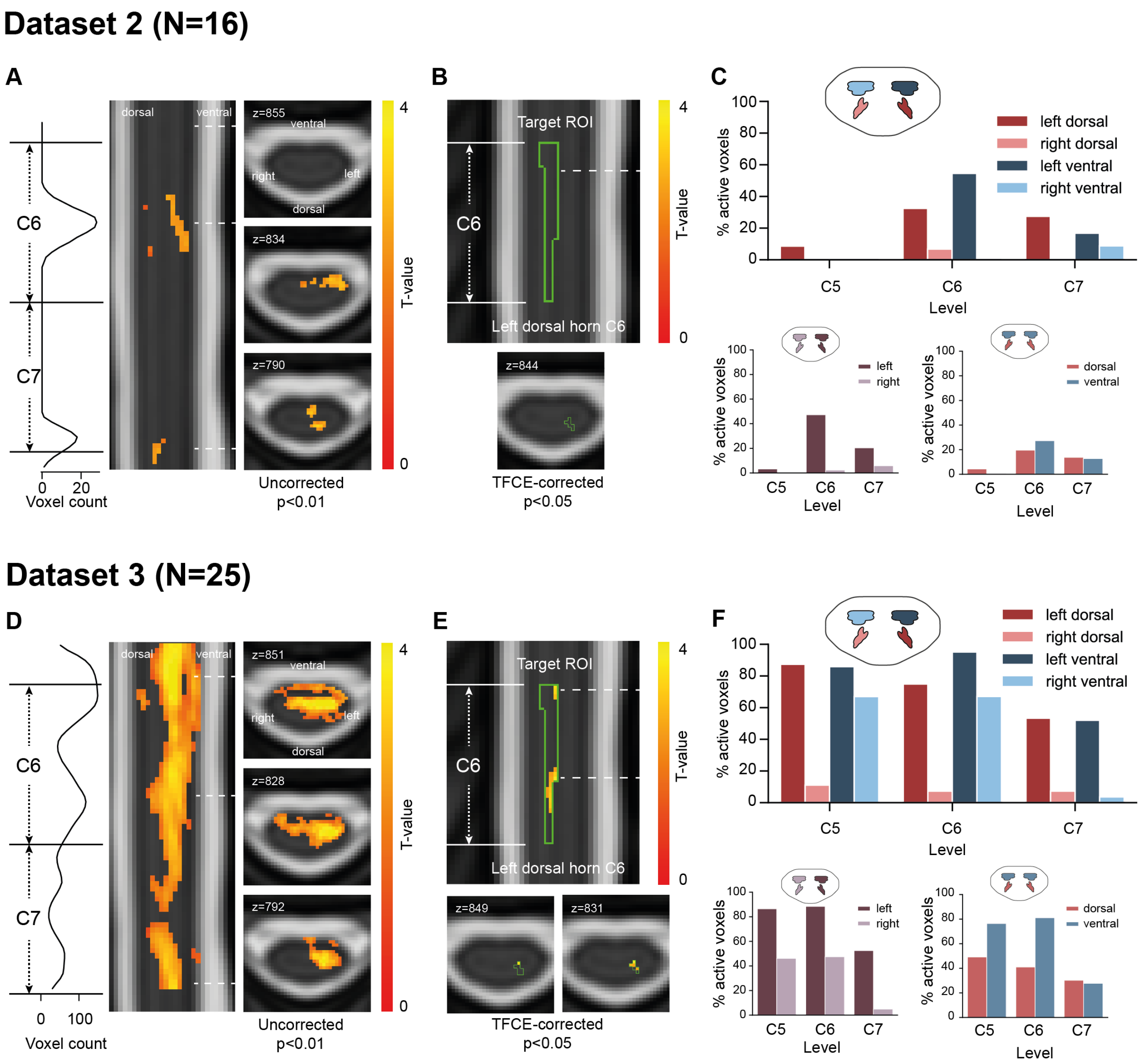


**Figure S1. Group-level spinal cord BOLD responses (sustained response) for individual datasets. A.** Uncorrected group-level results ($p<0.01$) in spinal cord for Dataset 2 in response to 30s heat pain. The density plot (left) shows the rostro-caudal activation distribution, with segments C6 and C7 marked. The sagittal view (middle) shows activation spread across spinal levels with the dashed lines indicating the position of the axial plots (right) at maximum (possibly subthreshold) activation in cord segments C5, C6, and C7. **B.** TCFE-corrected results ($p_{FWE}<0.05$) for Dataset 2 in response to 30s heat pain in the target ROI (C6 left dorsal horn, green outline), showing that no voxels survive strict multiple comparison correction. Dashed line indicating maximum (sub-threshold) activation in target ROI. **C.** Percentage of suprathreshold voxels (uncorrected $p<0.05$) in each grey matter horn mask (top) and separated into either left and right masks (bottom left) or dorsal and ventral masks (bottom right). **D. – F.** same as **A. – C.**, but for Dataset 3. Here the TCFE-corrected results depict two surviving clusters with maxima shown in the axial plots below.

**b. Phasic response**

In analogy to the analyses reported for the sustained response above, we tested whether we would be able to detect spinal cord BOLD responses at stimulus onset (modelled using the first 3 s) within each dataset (i.e., the *discovery* dataset [Dataset 2] and the *validation* dataset [Dataset 3]), using identical analysis parameters as in the combined dataset. At an exploratory threshold ($p < 0.01$), we observed sparse activation in both datasets (compared to the more extended sustained responses depicted in Figure S1). Both datasets exhibited a peak in target segment C6, that is found primarily on the ipsilateral (left) side and centred consistently more dorsally than the sustained response pattern in both datasets (Figure S2A, S2C; cf Figure S1A, S1C); a further peak in segment C7 emerged in both datasets. Although Dataset 2 allowed for the discovery of a phasic spinal cord BOLD response (and inspired the pre-registered analysis of the phasic response in Dataset 3), this response did not survive strict statistical thresholding using permutation testing in the target region of the left dorsal horn in segment C6 (Figure S2B; peak voxel: $t = 2.97$, $p_{FWE} = 0.107$). In the higher-powered Dataset 3 however, one dorsal horn cluster in C6 survived strict statistical thresholding (Figure S2D; cluster extent: 26 voxels, peak voxel: $t = 5.00$, $p_{FWE} = 0.0008$).

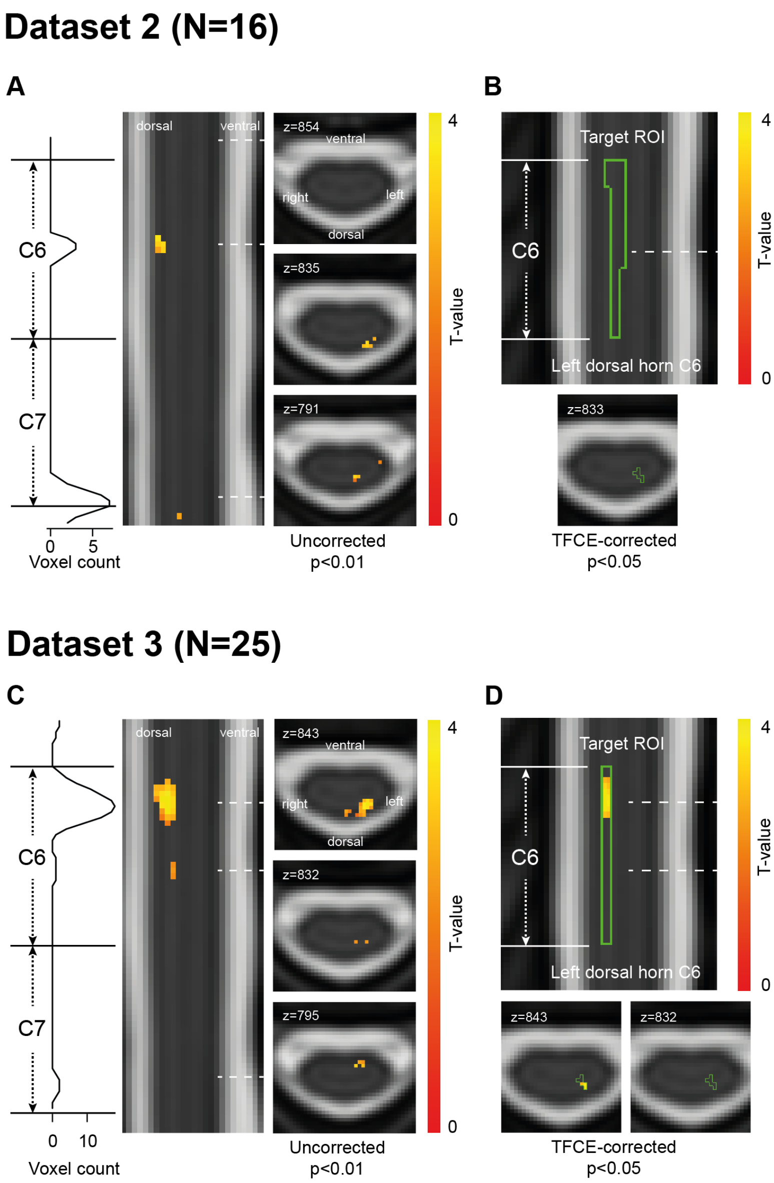


**Figure S2. Group-level spinal cord BOLD responses (phasic response) for individual datasets. A.** Uncorrected group-level results ($p<0.01$) constrained within the spinal cord for Dataset 2 in response to 3 s heat pain. The density plot (left) shows the rostro-caudal activation distribution, with segments C6 and C7 marked. The sagittal view (middle) shows sparse activation in C6 and C7 with the dashed lines indicating the position of the axial plots (right) at maximum (possibly subthreshold) activation in cord segments C5, C6, and C7. **B.** TCFE-corrected results ($p_{FWE}<0.05$) for Dataset 2 in response to 3 s heat pain in the target ROI (C6 left dorsal horn, green outline) showing that no voxels survive strict multiple comparison correction. Dashed line indicating maximum (sub-threshold) activation in target ROI. **C. & D.** same as **A. & B.**, but for Dataset 3. Here the dashed lines and axial plots depict maximum activation location of the two clusters in C6 of which only one survives strict multiple comparison correction.

### 3. Further autonomic and subjective heat pain responses

The behavioural dataset (Dataset 4) was used to assess whether sustained and phasic response components would also be evident in supraspinally mediated responses, i.e. in subjective and autonomic metrics of pain processing (Figure 5F-I). A subset of the autonomic measures was also acquired during fMRI acquisition (skin conductance data in Dataset 2 and heart period data in Datasets 2 and 3) and results from those acquisitions are depicted here. The response patterns appear almost identical to the ones acquired outside the MRI in Dataset 4 (cf. Figure S3A and Figure 5G, Figure S3B and Figure 5I).

Dataset 4 also allowed us to assess in detail the subjective percept resulting from sustained or phasic heat pain. More specifically, at the end of the experiment in Dataset 4, each participant received two more heat-pain stimuli: one with a duration of 30 s (i.e. identical to the stimuli used throughout the experiment in Datasets 2-4) and one with a duration of 3 s (i.e. identical to what was modelled using the 'phasic response' GLM in Datasets 2 & 3). We used the McGill Pain Questionnaire (MPQ; (Melzack, 1975); translated to German as in (Tal, 2008)) to allow participants to describe their pain percept to both types of stimuli. The group-level ratings on the MPQ descriptors show that the 30 s stimuli in general lead to a higher rating of several aspects of pain perception, but both stimuli (30 s and 3 s) are described using similar categories (e.g. hot-burning, aching, tender, stabbing, sharp; Figure S3C).

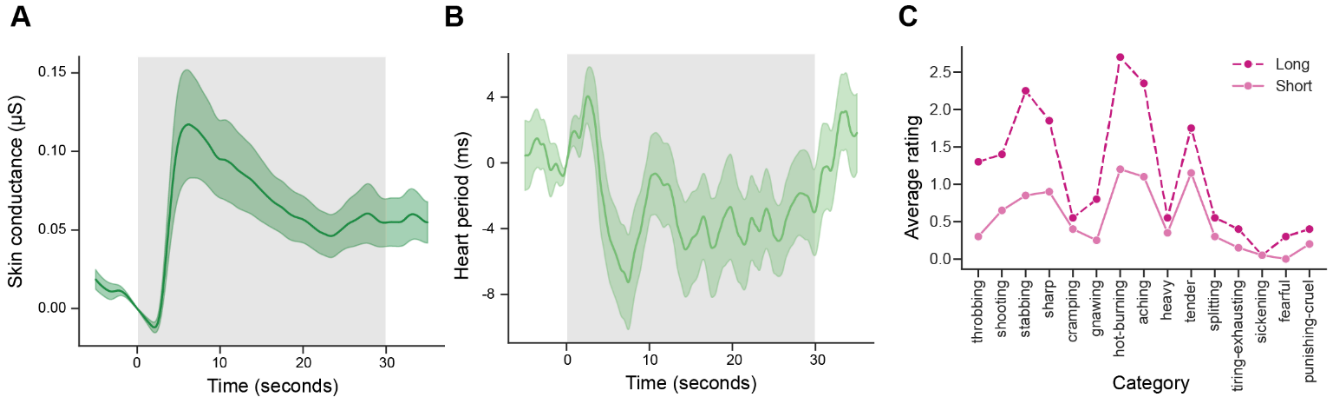


**Figure S3. Additional autonomic and subjective heat pain responses. A.** Averaged skin conductance in Dataset 2 (concurrent acquisition with 7T fMRI data). **B.** Averaged heart period responses across Datasets 2 and 3 (concurrent acquisition with 7T fMRI data). **C.** Group-level McGill Pain Questionnaire (MPQ) results in Dataset 4 in response to a 30 s heat stimulus (long) and a 3 s heat stimulus (short) of the same temperature.

### 4. Control analyses layer-specific spinal cord BOLD responses

#### a. Unsmoothed data

In addition to the layer-specific results presented in the main manuscript that are based on minimally-smoothed data (2 mm FWHM isotropic smoothing; i.e. analogous to all other analyses carried out), we here present layer-specific results from analyses on unsmoothed data. These analyses show virtually identical patterns with most frequent activation peaks in the deep layers for the sustained response (Figure S4B) and most frequent activation peaks in the superficial layers for the phasic response (Figure S4C). At the participant level, average individual beta values per layer were again tested in a one-tailed t-test against zero, revealing that for the sustained response only the superficial dorsal horn layer activation did not significantly differ from zero ($t(38) = 0.87$, $p = 0.20$, $d = 0.14$), with the deep dorsal and ventral horn showing the most prominent effects (middle dorsal horn: $t(38) = 3.24$, $p = 0.001$, $d = 0.52$; deep dorsal horn: $t(38) = 3.87$, $p = 0.0002$, $d = 0.62$; ventral horn: $t(38) = 4.37$, $p = 4.67 \cdot 10^{-5}$, $d= 0.70$). For the sustained response, the superficial dorsal horn activation was significantly weaker than the deep dorsal horn activation ($t(38) = -2.49$, $p = 0.017$, $d = -0.40$). Parallel to the minimally smoothed data, the analyses for the unsmoothed data reveal a similar dominance of superficial dorsal horn responses in the phasic response analysis (superficial dorsal horn: $t(38) = 2.50$, $p = 0.009$, $d = 0.40$; middle dorsal horn: $t(38) = 0.32$, $p = 0.37$, $d = 0.05$; deep dorsal horn: $t(38) = 0.12$, $p = 0.45$, $d = 0.02$; ventral horn: $t(38) = 0.89$, $p = 0.19$, $d = 0.14$). Only for the phasic response, we could not establish significantly stronger activation in the superficial dorsal horn activation compared to the deep dorsal horn ($t(38) = 1.64$, $p = 0.11$, $d = 0.26$); note that this is a two-tailed test as this analysis was not preregistered.

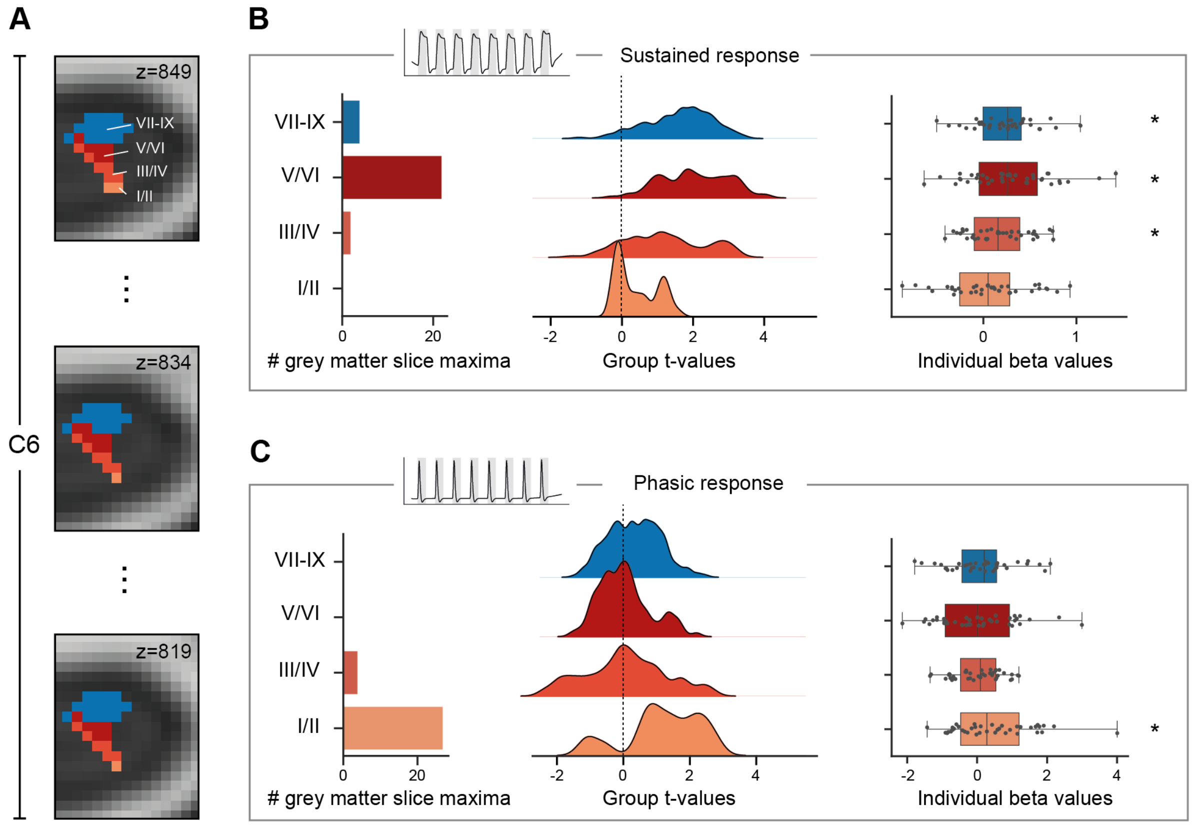


**Figure S4. Layer-specific spinal cord BOLD responses in unsmoothed data. A.** A newly-created and histology-based atlas in PAM50 template space distinguishes layers within the grey matter (depicted is the left hemicord): superficial dorsal horn (laminae I/II), middle dorsal horn (laminae III/IV), deep dorsal horn (laminae V/VI), and ventral horn (laminae VII-IX); exemplary C6 slices are displayed. **B.** Layer-specific BOLD responses for the sustained heat GLM: slice-wise occurrence of the group result's maximum grey matter voxel in each layer (left); distribution of group analysis t-values in each layer (middle); individual participants' average activation (beta) values in each layer (right; asterisk marks significance for test against 0). **C.** Layer-specific BOLD responses for the phasic heat GLM, with panels organized as in B.

### b. Individual datasets

In addition to the layer-specific results presented in the main manuscript and based on the combined dataset, we here present results from Dataset 2 and Dataset 3 separately.

*Sustained response*

Similar to the combined result, in Dataset 2 the one-tailed t-test of individual beta values per layer revealed that for the sustained response only the superficial dorsal horn layer activation did not significantly differ from zero ($t(15) = -0.65$, $p = 0.74$, d = -

0.16), with the deep dorsal and ventral horn showing the most prominent effects (middle dorsal horn: $t(15) = 2.04$, $p = 0.029$, $d = 0.51$; deep dorsal horn: $t(15) = 2.20$, $p = 0.022$, $d = 0.55$; ventral horn: $t(15) = 2.22$, $p = 0.021$, $d= 0.56$). In parallel to the combined results, the superficial dorsal horn activation was significantly weaker than the deep dorsal horn activation ($t(15) = -2.32$, $p = 0.035$, $d = -0.58$).

In Dataset 3, the t-test against 0 revealed that all regions were significantly activated, even though the superficial dorsal horn showed by far the weakest effect (superficial dorsal horn: $t(24) = 2.03$, $p = 0.027$, $d = 0.41$; middle dorsal horn: $t(24) = 4.43$, $p = 0.000088$, $d = 0.89$; deep dorsal horn: $t(24) = 5.38$, $p = 0.000008$, $d = 1.07$; ventral horn: $t(24) = 5.38$, $p = 0.000023$, $d= 0.99$) and consequently, there was again a significant difference between superficial and deep dorsal horn ($t(24) = -2.59$, $p = 0.016$, $d = -0.52$).

*Phasic response*

In Dataset 2, the results for the phasic response did not match the combined results as the beta values did not significantly differ from 0, though the pattern of results was similar with the superficial horn showing the strongest response (superficial dorsal horn: $t(15) = 0.89$, $p = 0.19$, $d = 0.22$; middle dorsal horn: $t(15) = -0.20$, $p = 0.577$, $d = -0.05$; deep dorsal horn: $t(15) = -0.59$, $p = 0.72$, $d = -0.15$; ventral horn: $t(15) = 0.59$, $p = 0.28$, $d = 0.15$). The test for a difference between superficial and deep dorsal horn responses was not significant ($t(15) = 1.52$, $p = 0.15$, $d = 0.38$).

In Dataset 3, the results for the phasic response are similar to the combined results revealing a similar dominance of superficial dorsal horn responses in the phasic response analysis (superficial dorsal horn: $t(24) = 4.09$, $p = 0.00021$, $d = 0.82$; middle dorsal horn: $t(24) = 1.39$, $p = 0.089$, $d = 0.28$; deep dorsal horn: $t(24) = 0.42$, $p = 0.34$, $d = 0.08$; ventral horn: $t(24) = 0.23$, $p = 0.41$, $d = 0.04$). Similar to the combined results, the superficial dorsal horn activation was significantly stronger than the deep dorsal horn activation ($t(24) = 2.17$, $p = 0.039$, $d = 0.43$; two-tailed).

#### c. Voxel-wise analyses

In addition to the tests involving individual participants' average activation (beta) values in each layer, we also conducted voxel-wise t-tests via non-parametric permutation testing as implemented in FSL's randomise algorithm (Winkler et al., 2014) using Threshold-Free Cluster Enhancement (Smith & Nichols, 2009) and controlling the family-wise error rate (FWE). The results for these analyses match the ones obtained using participants' individual beta values.

For the sustained response, all atlas regions except the superficial dorsal horn are significantly activated (superficial dorsal horn: no cluster, p=0.195; middle dorsal horn: 89 voxels, p=0.0006; deep dorsal horn: 186 voxels, p=0.0002; ventral horn: 358 voxels, p=0.0006). After applying a Bonferroni correction for multiple comparisons, the significance level was adjusted to 0.0125. All the reported p-values are below this adjusted threshold.

For the phasic response, only the superficial dorsal horn activation survives this stringent correction, which fits the results obtained using participants' individual beta values (superficial dorsal horn: 10 voxels, p=0.001; middle dorsal horn: 1 voxel, p=0.0468; deep dorsal horn: no cluster, p=0.643; ventral horn: no cluster, p=0.643).

### 5. Differences between preregistration and manuscript

**Table S1. Differences between preregistration and manuscript.** Listed are all differences between what was specified in the preregistration and what is reported in the manuscript.

| **Preregistration** | **Manuscript** |
|---|---|
| EPI flip angle: 61°. | EPI flip angle ideally set to 61°, but if individually optimal reference voltage exceeded the maximum allowable for the coil, the reference voltage was set to the maximum value and the flip angle was adapted accordingly. |
| Target sample size Dataset 3: 17. | Final sample size Dataset 3: 25 due to availability of additional resources (see section 2.1 for further motivation). |
| Slice-wise motion correction in two steps. | Slice-wise motion correction in two steps was complemented by a novel procedure for through-slice motion correction. |
| Analysis of fMRI data using GLM modelling whole 30 s and GLM modelling first 3 s. | Analysis of fMRI data using GLM modelling whole 30 s and GLM modelling first 3s was complemented by an additional GLM |

| | |
|---|---|
| | comparing even and odd trials to investigate reliability. |
| Spatial smoothing along the spinal cord centerline with a 2×2×6 mm kernel + reporting results with and without smoothing. | Spatial smoothing within FSL FEAT with a 2 mm isotropic kernel to allow for layer-specific analyses, which are reported with and without smoothing. |
| Additional analyses to account for larger spatial spread will be carried out for the remaining levels of our slice stack. | For the sake of brevity, we refrained from also conducting ROI analyses in other spinal segments, but show uncorrected results for the whole slice stack and individual analyses focusing on the spread of responses. |
| | Although we did not preregister the VasA method (the use of which arose from separate work in our lab), its implementation reveals numerous advantages without any apparent disadvantages. |
